# Energetics and kinetics of membrane permeation of photoresists for bioprinting

**DOI:** 10.1101/2023.03.27.534360

**Authors:** Lucas Diedrich, Matthias Brosz, Tobias Abele, Salome Steinke, Frauke Gräter, Kerstin Göpfrich, Camilo Aponte-Santamaría

## Abstract

Three-dimensional (3D) bioprinting is a promising technology which typically uses bioinks to pattern cells and their scaffolds. The selection of cytocompatible inks is critical for the printing success. In laser-based 3D bioprinting, photoresist molecules are used as bioinks. We propose that cytotoxicity can be a consequence of the interaction of photoresists with lipid membranes and their permeation into the cell. Here, molecular dynamics simulations and in vitro assays address this issue, retrieving partition coefficients, free energies, and permeabilities for eight commonly-used photoresists in model lipid bilayers. Crossing the hydrophobic center of the membrane constitutes the rate limiting step during permeation. In addition, three photoresists feature a preferential localization site at the acyl chain headgroup interface. Photoresist permeabilities range over eight orders of magnitude, with some molecules being membrane-permeable on bioprinting timescales. Moreover, permeation correlates well with the oil-water partition coefficients and is severely hampered by the lipid ordering imposed by the lipid saturation. Overall, the mechanism of interaction of photoresists with model lipid bilayers is provided here, helping to classify them according to their residence in the membrane and permeation through it. This is useful information to guide the selection of cytocompatible photoresists for 3D bioprinting.

## 1 Introduction

3-dimensional (3D) bioprinting utilizes 3D printing technologies to manipulate and arrange cells and their scaffolds. Applications range from single cell research [1] to the fabrication of biomedical parts which imitate natural tissues and organs [2]. Various 3D printing techniques have been developed for this purpose, such as inkjet printing [3], extrusion printing [4], or direct laser writing [5], which vary tremendously in printing rate and resolution. Extrusion-based printing techniques have been used to provide scaffolds for 3D cell culture on a macroscale [6]. Moreover, larger photopatterned hydrogel geometries have been used to provide extracellular cues for organoid morphogenesis [7]. On the micro scale, single cells have been seeded on scaffolds produced by two-photon direct laser printing to gain fundamental insights on single cell adhesion [8]. Functional micro-scaffolds also allowed to interact with cells live during an ongoing experiment, e.g. to exert mechanical stress [9]. Moreover, it has been possible to print the cells themselves using cell-laden photoresists to generate tissue-like arrangements [10]. When cells are present during the printing process or when they are printed themselves, photoresists, i.e. the molecules used during laser-based 3D bioprinting, have to be cytocompatible.

The cell membrane provides a semi-permeable barrier that tightly regulates the passage of solutes in and out of the cell. Membranes establish and maintain concentration gradients of protons, ions, metabolites or toxic waste products and thus sustain life as an out-of-equilibrium process. In bioprinting experiments, it is commonly assumed and expected that the bioinks do not enter the cell. However, recent experiments with giant unilamellar lipid vesicles (GUVs) have shown that commonly used photoresists, including photoinitiators and monomers, can pass across the membrane by passive diffusion [11]. This feature has been exploited to print micrometer-scale 3D objects of arbitrary shape inside of synthetic cells using twophoton laser printing [11]. Permeation of resist molecules across cellular membranes may have complex downstream effects that could contribute to cytotoxicity. To overcome cytotoxicity of photoresists and to thereby advance on the applicability of laser-based bioprinting, it is crucial to understand how commonly used photoresists interact and pass the membrane.

Molecular dynamics (MD) simulations can provide insights into the process of membrane permeation at a molecular level of resolution. MD simulations have been widely employed to elucidate the molecular principles governing membrane permeation and to quantitatively predict membrane permeabilities for a broad range of substances including oxygen and water [12], simple organic molecules like alcohols [13], druglike molecules [14, 15, 16] or fluorescent dyes [17]. In contrast to other approaches, such as quantitative structure-permeability relationship models that predict membrane permeability from available physicochemical descriptors of molecules [18], MD simulations further provide detailed information about the energetics, local kinetics, and molecular interactions. This information has been further used to delineate the principles behind lipid-molecule interactions [12, 14, 19] and to inform experiments [17]. Hence, MD simulations are ideally suited, in particular in conjunction with experiments, to guide the right choice of photoresists for bioprinting applications.

Here, we combine experiments and MD simulations to provide the molecular basis of the—so far poorly understood—interplay between photoresists and biological lipid bilayers. Using partitioning and influx experiments as well as non-equilibrium and biased MD simulations, we explain (i) the thermodynamic propensity of the photoresist molecules to reside in the oil phase of the lipid bilayer and (ii) their kinetic ability to permeate through lipid bilayers of distinct lipid composition. Ultimately, we show that our computational workflow can provide valuable insights into the permeation of photoresists across biological membranes complementing the experimental data. This provides a possible way to classify photoresists in terms of their interaction with cells and their potential to exert cytotoxicity. All in all, we thus provide a framework to design and select new biocompatible resists for 3D bioprinting.

## 2 Results and Discussion

### 2.1 Choice of photoresists for membrane permeation studies

Figure 1A illustrates an experiment to assess whether a photoresist is membrane permeable. We immerse cell-sized giant unilamellar lipid vesicles (GUVs) in a photoresist solution in an attempt to perform direct 3D laser printing outside as well as inside the GUV. If printing is possible inside the GUV, the photoresist molecules must have passed across the lipid membrane by passive diffusion along the concentration gradient. Computationally, we reproduce this setup by placing photoresists in a minimalistic representation of the membrane system, consisting of 162 lipid molecules (Figure 1B).

**Figure 1:**
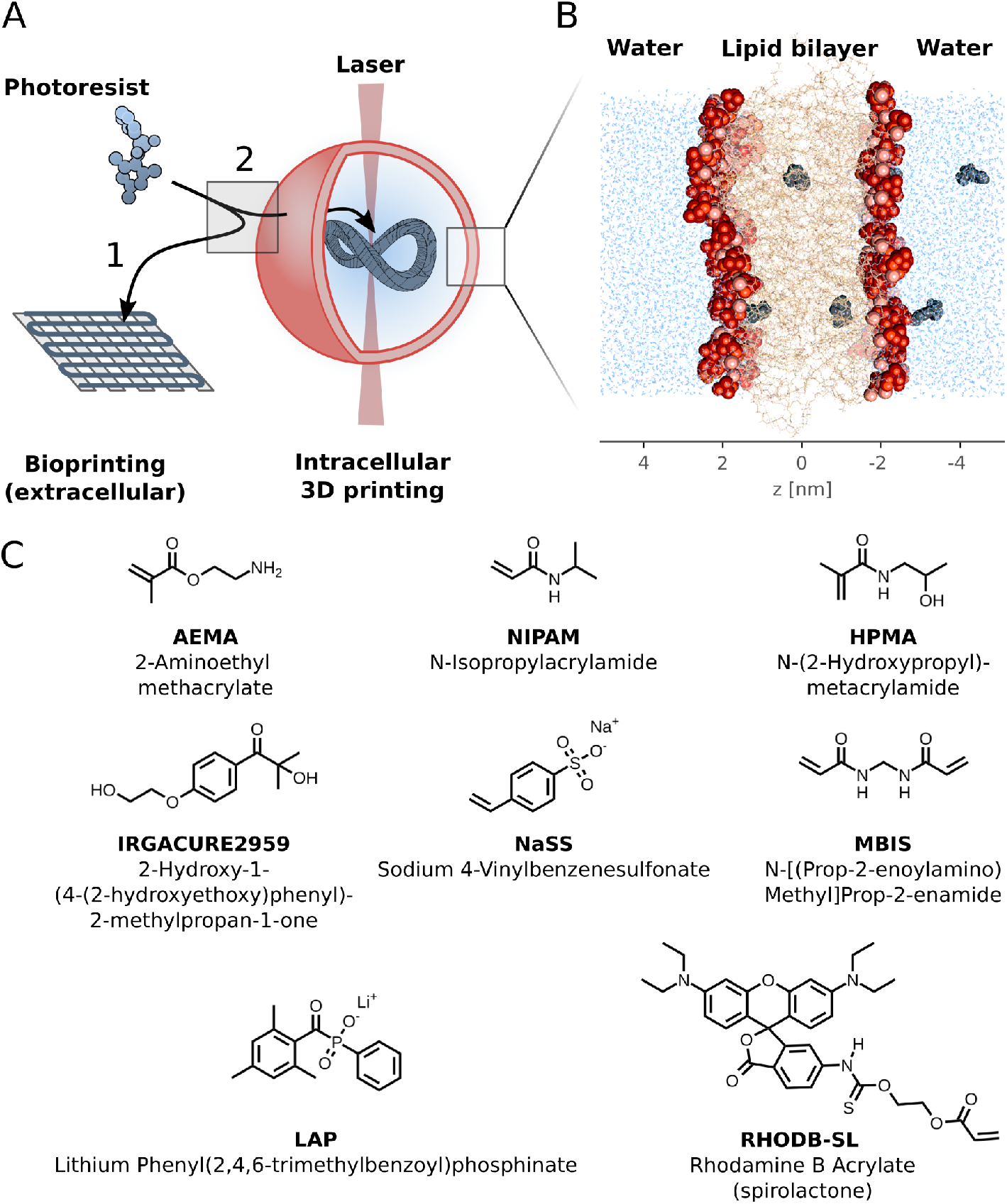
3D bioprinting at the interface of biological lipid bilayers. A) The photoresists necessary for 3D printing, either near (1) or inside (2) biological compartments enclosed by a lipid bilayer (red) requires spatio-speficic polymerization of photoresist monomers, for instance by two-photon laser printing [11]. In scenario 2, photoresists permeate the lipid bilayer by passive diffusion. B) A fragment of the surrounding membrane constituted by a lipid bilayer (here of 1-palmitoyl-2-oleoylsn-glycero-3-phosphocholine, POPC lipids). Lipid headgroups are shown as red spheres, phosphor atoms as light red spheres, and acyl chains as orange sticks. Bulk water is colored blue. Ions are not shown. Photoresist molecules which cross the bilayer are depicted in dark gray. To investigate the energetics and permeation of such molecules MD simulations were carried out by placing several molecules in two columns in the membrane system. C) Shown are the molecular structures, names and, if applicable, abbreviated names of the photoresist molecules whose permeation through lipid bilayers was examined in this study.

We selected a representative set of eight photoresist molecules which are commonly used in 3D bioprinting [20], namely AEMA, NIPAM, HPMA, Irgacure 2959, NaSS, MBIS, LAP and RHODB-SL (Figure 1C). With this choice of molecules, we aimed at covering distinct physico-chemical properties commonly related to their interaction with biological membranes. These properties include molecular weight, hydrophobicity, number of rotatable bonds, number of hydrogen bond donors, and number of hydrogen bond acceptors [21, 22, 23] (Figure S1). Notably, two of the examined photoresist molecules, i.e. LAP and NaSS, contain a negative charge, which is neutralized by lithium and sodium, respectively.

### 2.2 Classification of resists according to their oil-water partition coefficients

We investigated the oil-water partitioning of the studied photoresist molecules both computationally (Figure 2A) and experimentally (Figure 2B), by determining octanol-water partition values (log *K*_*OW*_). This quantity provides insights into the thermodynamic preferential location of the resists in oil-water biphasic systems, such as that present near water-membrane interfaces. Since the partition coefficient is a quantity that is accessible both experimentally and computationally, it can also be used to validate the obtained force field parameters for the photoresist molecules employed in the simulations.

**Figure 2:**
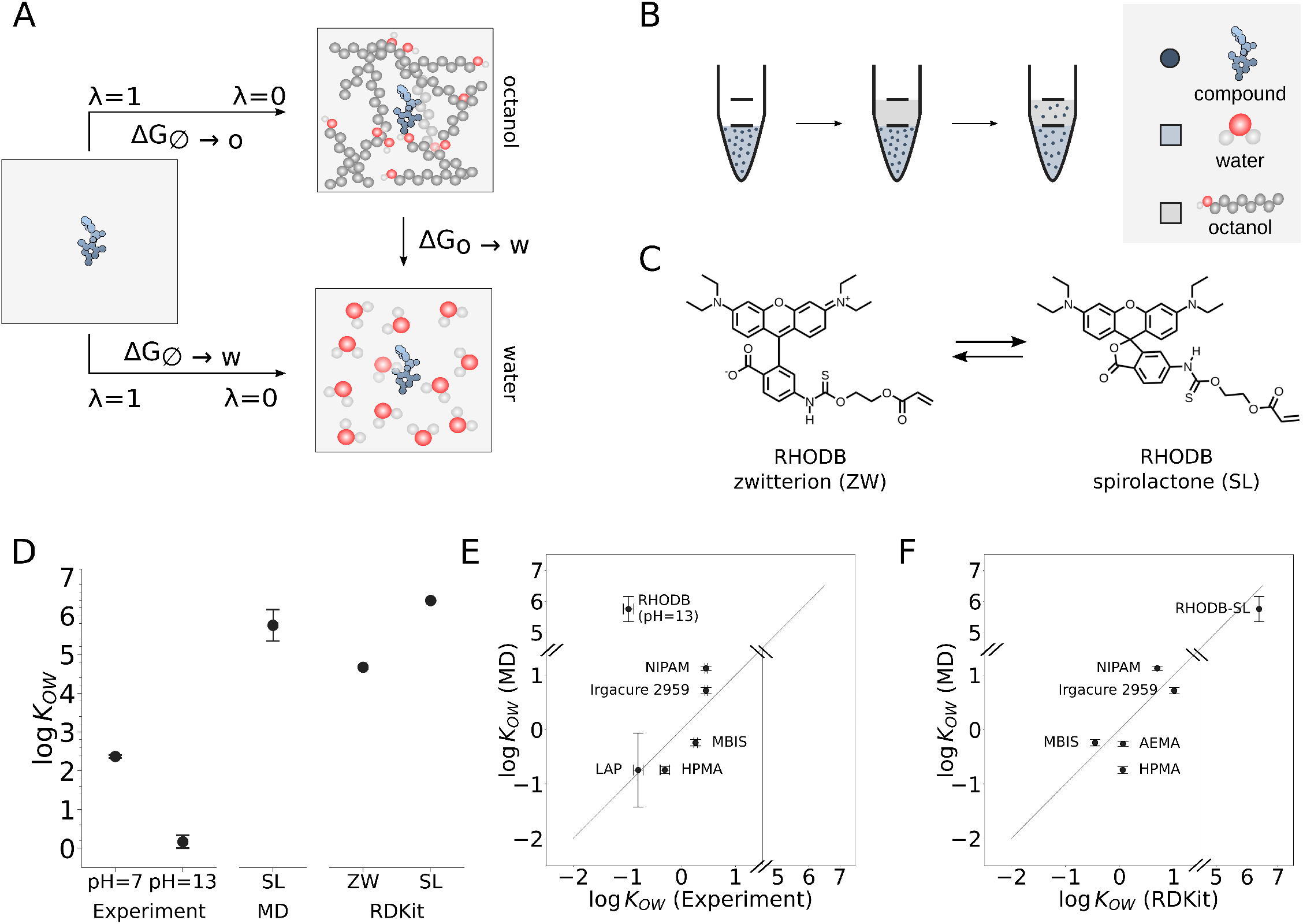
Determination of log *K*_*OW*_ values with MD simulations and experiments. A) Computation of log *K*_*OW*_ values with non-equilibrium MD-based free energy calculations. The log *K*_*OW*_ is related to the free energy of transfer from octanol into water (Δ*G*_*o*→*w*_). This quantity was determined by separately computing the free energy of transfer the photoresist molecule (blue molecule) from vacuum into octanol (Δ*G*_∅→*o*_) or into water (Δ*G*_∅→*w*_), i.e. the solvation free energies in these two solvents, according to eq. 1. The molecule was made to “disappear” from the solvent in multiple non-equilibrium simulation runs, by gradually switching off its interaction with the solvent varying the *λ* parameter from 0 to 1. B) Experimental determination of the log *K*_*OW*_ value. The compounds were dissolved in water. The water phase was thoroughly mixed with the apolar octanol phase until an equilibrium was reached and the concentration of the resist was subsequently measured in the water phase using absorbance or osmolality measurements. The obtained concentrations were used to determine the log *K*_*OW*_ value according to eq. 8. C) Equilibrium between the zwitterionic, fluorescent form RHODBZW and the spirolactone form RHODB-SL with its closed ring conformation. D) log *K*_*OW*_ value for RHODB-SL determined experimentally at the indicated pH values, obtained from MD simulations (using the protocol described in A), and predicted by the cheminformatics machine-learning tool RDKit. For the computational predictions, either the spirolactone (SL) or the zwitterionic (ZW) forms of the molecule were used. E) Correlation between experimental and computed log *K*_*OW*_ values (without RHODB-SL *r* = 0.84 and RMSE = 0.44 were obtained). The gray line indicates ideal correlation. F) Correlation between log *K*_*OW*_ values computed from MD simulations and by RDKit (*r* = 0.98, RMSE = 0.50). The gray line indicates ideal correlation. Uncertainties in the values from the simulations were estimated by bootstrapping and error propagation (see methods). Experimental values correspond to averages ± standard deviation.

In our computational approach, we performed non-equilibrium MD-based free energy calculations to determine the solvation free energy of the photoresist molecules in water and octanol, separately, and subsequently used these energies to obtain the log *K*_*OW*_ value by means of the thermodynamic cycle shown in Figure 2A. We also predicted log *K*_*OW*_ with the cheminformatics tool RDKit [24], which uses an orthogonal, atom-typing based machine-learning approach [25], to predict octanol-water partition coefficients.

Experimentally, we obtained octanol-water partition coefficients by layering octanol on top of the aqueous phase containing the photoresist molecule of interest (Figure 2B). After thorough mixing to enable partitioning, we separated the phases via centrifugation, extracted the aqueous phase and compared the concentration before and after the partitioning experiment by measuring the absorbance or osmolality of the solution (Figure 2B). The obtained log *K*_*OW*_ values for the three approaches are listed in Table 1.

**Table 1:** Partition coefficient for each photoresist. Comparison of experimental log *K*_*OW*_ values, computed log *K*_*OW*_ values from MD simulations and predictions with RDKit

| Name | Experiment | MD | RDKit |
| --- | --- | --- | --- |
| AEMA | NA | $-0.26 \pm 0.05$ | 0.06 |
| HPMA | $-0.31 \pm 0.09$ | $-0.74 \pm 0.07$ | 0.06 |
| Irgacure 2959 | $0.46 \pm 0.02$ | $0.72 \pm 0.06$ | 1.01 |
| LAP | $-0.80 \pm 0.09$ | $-0.74 \pm 0.68$ | NA |
| MBIS | $0.26 \pm 0.04$ | $-0.24 \pm 0.06$ | -0.45 |
| NaSS | NA | $-6.35 \pm 0.70$ | NA |
| NIPAM | $0.46 \pm 0.02$ | $1.13 \pm 0.04$ | 0.70 |
| Rhodamin B Acrylate (zwitterion) | $2.37 \pm 0.04$ | NA | 4.67 |
| Rhodamin B Acrylate (pH=13) | $0.17 \pm 0.17$ | NA | NA |
| Rhodamin B Acrylate (spirolactone) | NA | $5.75 \pm 0.40$ | 6.39 |

For several photoresist molecules, the experimental determination of log *K*_*OW*_ turned out to be challenging. On the one hand, the concentration of both AEMA and NaSS could not be measured spectroscopically, due to a low extinction coefficient (in the case of AEMA) or presumably due to its charge (in the case of NaSS). Furthermore, the charged state of the molecules in neutral solution (NaSS: negatively charged sulfo group, AEMA: positively charged amino group) prohibited the determination of their concentration via solution’s osmolality measurements.

For the fluorescent dye Rhodamine B acrylate (RHODB), on the other hand, it has been established that the molecule occurs in an equilibrium between a rather polar, highly abundant zwitterionic form (RHODBZW) and an energetically unfavorable, uncharged spirolactone (RHODB-SL) form (Figure 2C). The latter is the primary membrane permeable form [26, 27] and was therefore considered in this study. Previous reports for comparable Rhodamine dyes suggested that the equilibrium between zwitterionic and spiro form shifts towards the latter at high pH values [28, 29, 30]. Following these studies, we attempted to determine the partition coefficient for this compound at pH=13. The obtained log *K*_*OW*_ values for this molecule are shown in Figure 2D. Surprisingly, the measured partition coefficient at high pH was smaller than the one at neutral pH values, contrary to the chemical intuition that RHODB-SL should be more hydrophobic than RHODB-ZW. In fact, both simulation and cheminformatics computational predictions predicted much higher values for RHODB-SL. Moreover, we observed that even at high pH values, the absorbance of the dye did not show a significant decrease, as previously observed for comparable Rhodamine-derived molecules [28, 29, 30] (Figure S3). This suggests that the zwitterionic species, i.e. the one typically found at neutral pH, was still present in solution. Accordingly, the reported log *K*_*OW*_ values most likely represent the octanol-water partition coefficient of the zwitterionic form. Experimentally establishing a log *K*_*OW*_ value for the membrane permeable form of Rhodamine B Acrylate, RHODB-SL, remains challenging with the assays we employed. Therefore, the obtained values rather provide a lower boundary of the log *K*_*OW*_ value expected for RHODB-SL.

Where applicable, we compared our computational and experimental predictions for the log *K*_*OW*_ value and found them to be in good agreement, with a root mean squared error RMSE = 0.38 and a Pearson correlation coefficient *r* = 0.84, considering all compounds except the challenging RHODB-SL (Figure 2E). The experimental log *K*_*OW*_ values span a wide range between approximately −0.8, for the rather hydrophilic initiator LAP, and around 0.5, for the more hydrophobic NIPAM or Irgacure 2959 molecules. To further validate the parametrization of the other photoresists, we compared our molecular dynamics-derived log *K*_*OW*_ -predictions with the log *K*_*OW*_ -predictions of the cheminformatics tool RDKit [24, 25]. For RHODB-SL and AEMA, as well as the experimentally accessible photoresist molecules, both computational methods agree well (RMSE = 0.50 a.u., Pearson *r* = 0.98, Figure 2F). In particular, the prediction of both computational approaches suggests RHODB-SL to be very hydrophobic, a finding that is consistent with the notion that it is the membrane permeable form [26, 27].

Finally, for NaSS, MD simulations predict a log *K*_*OW*_ value of −6.35 ± 0.70, indicating this to be a very hydrophilic molecule. For this molecule the log *K*_*OW*_ could not be obtained, neither experimentally (see above) nor with RDKit, which is only valid for uncharged molecules [25, 31]. Precisely this molecule’s dominant hydrophilic character results in a very large free energy barrier of membrane crossing as shown in the next section.

Altogether, our comparison between simulation, cheminformatics, and experimental log *K*_*OW*_ data validates the chosen force-field parameters for the studied set of photoresists. Beyond that, we quantitatively classify these resist molecules according to their propensity to locate in rather hydrophobic solvents (RHODB-SL, NIPAM or Irgacure 2959) versus water (e.g. LAP, NaSS). This is important thermodynamic information for the intended use of these molecules in 3D bioprinting near or inside biological substrates. Moreover, our results underline how molecular dynamics can complement experiments by providing physico-chemical properties of molecules which are not always accessible experimentally.

### 2.3 Computation of free energy profiles reveal preferential locations of photoresists in lipid bilayers

To gain further insights into the interplay between photoresist molecules and biological lipid bilayers, we computed free energy profiles, i.e. the potential of mean force of the photoresist as a function of its position perpendicular to the membrane, taking the bulk water as reference (Figure 3A). To this end, we used umbrella sampling MD simulations as a method to enhance the sampling. In a minimalistic representation of the compartmentalizing membrane, we considered a pure lipid bilayer of 162 1-Palmitoyl-2-oleoylphosphatidylcholine (POPC) lipids. Phosphorous–phosphorous distance (thickness), area per lipid, and deuterium order parameter, recovered from equilibrium MD simulations for this bilayer, were found to be in good agreement with previous experimental estimates, thus validating the chosen simulation protocol for this type of lipids (Table S3 and Figure S4). We placed multiple molecules of one photoresist species simultaneously to further enhance sampling as previously suggested in ref. [32] (Figure 1B).

**Figure 3:**
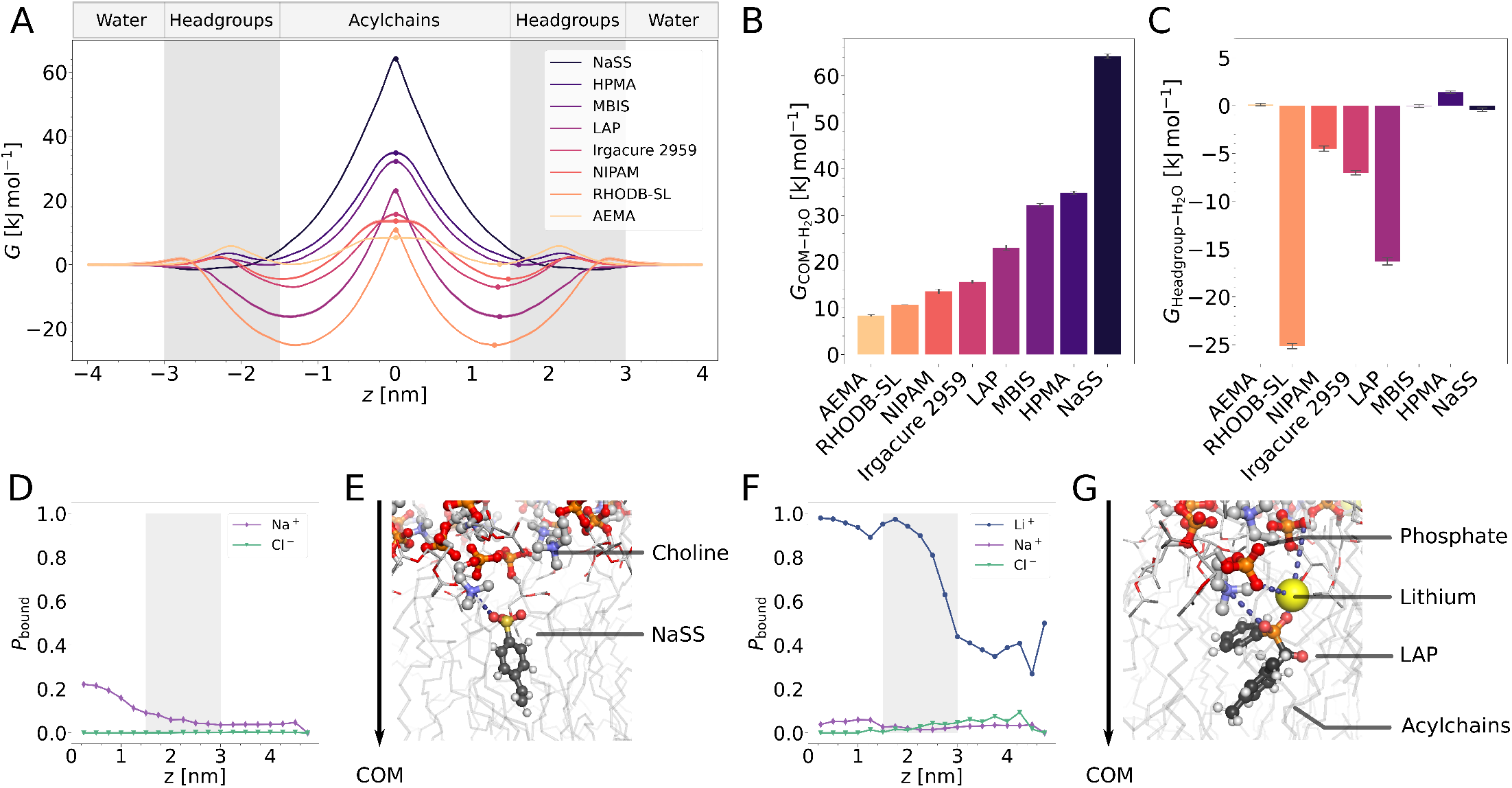
Energetics and kinetics of permeation of photoresists across a POPC lipid bilayer. A) Free energy profile, *G*, as a function of the position of the photoresist orthogonal to the membrane *z*. The profiles are mirrored at the center of the membrane (*z* = 0 nm). The displayed profiles are based on the aggregated analysis of three independent replicates. The approximate location of bulk water, headgroups and hydrophobic acyl chains regions is indicated. *G* is given relative to the water bulk media (i.e. *G* for the water media equals 0). B–C) Extracted *G* values at characteristic points of the profile and their bootstrapping errors at the center of membrane (COM) (B) and headgroup (C) positions, relative to the water media. D–G) Differences in membrane-molecule interactions for the ionic molecules NaSS and LAP. D,F) Probability of observing an ion bound to the photoresist molecule, *P*_*bound*_, between the ionic molecule and free ions (Li^+^, Na^+^, Cl^-^) as a function of the permeation coordinate *z* for NaSS (D) and LAP (F). A binding event was observed when the distance between an ion and the ionic part of the molecule is less than 0.5 nm. E,G) Representative snapshots of the interaction of NaSS and the membrane lipids in the center of membrane (*z* = 0.08 nm, E) and the binding of lithium to LAP when LAP is located in the hydrophobic center of membrane (*z* = 0.12 nm, *t* = 50 ns, G). The arrow points towards the center of the membrane.

Although the free energy profiles varied among photoresist molecules, they all displayed an energy barrier at the hydrophobic center of the membrane (Figure 3A,B). For many resists, mainly the more hydrophilic ones, a second, smaller energy barrier was observed at the headgroup region of the membrane, raising up to 6 kJ mol^−1^ for AEMA (Figure 3A). We attribute this barrier to the energetically unfavorable loss of the hydration shell upon entry into the headgroup region of the membrane.

The height of the central main barrier varied significantly between the different species (Figure 3A,B). Not surprisingly, we observe the greatest difference in the free energy between the center of membrane (COM) and bulk water for the ionic monomer NaSS (*G*_COM_ − *G*_H O_ = 64.2 ± 0.5 kJ mol^−1^). For this monomer, the sodium ion spontaneously dissociated from the negatively-charged sulfonate group causing a very unfavorable interaction between the resist molecule and the non-polar membrane center in comparison to its interaction with the polar water molecules (Figure 3D,E). Interestingly, for the other ionic molecule, LAP, the computed free energy difference is considerably lower (*G*_COM_ − *G*_H O_ = 23.0 ± 0.6 kJ mol^−1^). This may be attributed to the fact that the lithium ion stayed bound to the phosphinate group maintaining its neutrality and helping to bridge the phosphate groups of the lipids with negative charge in LAP while the hydrophobic aromatic rings were embedded in the acyl chains (Figure 3F,G). We observe this feature even at positions close to the COM suggesting that this headgroup-molecule interaction might help to reduce the energy barrier at the center of the acyl chains too. NaSS is lacking this strong interaction between its charged moiety, a positively charged ion and the lipid phosphogroups, presumably due to the lower charge density of the sulfonate- compared to the phosphinate group. The reduced barrier also aligns with the trend in our log *K*_*OW*_ calculations where we obtained a considerably higher value for LAP than for NaSS and predicted a strong accumulation of NaSS in the water phase compared to the organic phase (Table 1). Overcoming the main energetic barrier imposed by the hydrophobic acyl chains at the COM hence represents the rate-limiting step for the permeation of these molecules. Thus, we expect resists with a comparatively low barrier (e.g. AEMA, RHODB-SL, NIPAM) to permeate faster through the membrane than molecules with a high barrier, such as HPMA or NaSS (Figure 3B).

Several molecules, e.g. the photoinitiator LAP, the monomer NIPAM, or the fluorescent dye RHODB-SL, further exhibit a prominent energy well close to the headgroups of the membrane around 1.3 nm from the COM (Figure 3C). This indicates that the resists preferentially accumulate on the surface of the lipid bilayer. Conversely, this energy well was not observed for other molecules like AEMA or MBIS. LAP and RHODB-SL contain both hydrophobic aromatic groups and hydrophilic moieties and therefore the acyl chain-headgroup interface appears to be an energetically favorable location for them (Figure 3C). Interestingly, for LAP this favourable position occurred at the expense of the deformation of the membrane, bringing the phosphate lipid headgroups close to the phospinate group of LAP, bridged by a lithium ion (Figure 3G).

The obtained energetics quantify the propensity of the photoresist molecules to reside on the lipid membrane surface or instead in its aqueous surrounding. NaSS and HPMA may be considered membrane impermeable on experimentally relevant timescales, since we computed an energy barrier multiple times the thermal energy (≈ 2.54 kJ mol^−1^ at 310 K). On the contrary, NIPAM, LAP and RHODB-SL accumulate at the water-membrane interface and need to overcome comparably small barriers to permeate the membrane. In this way, our data can inform the choice of photoresist for bioprinting applications, providing an insight into how energetically favorable it is for these molecules to cross the lipid bilayer or to accumulate at the membrane, both of them potentially leading to undesired toxic effects.

### 2.4 From thermodynamics to permeabilities

A key question is to elucidate molecular principles explaining the cytotoxicity of bioinks used in 3D bioprinting. The computed energetics already gave a hint on the degree of membrane permeability of the resists. We next aimed at connecting this thermodynamic information with a kinetic descriptor, namely the membrane permeability of the photoresists. This quantity proved to be difficult to be determined experimentally, since a direct measurement of the concentration of a photoresist in a vesicle or cell is usually not possible, making standard kinetic measurements unfeasible. On the contrary, the free energy profiles, together with diffusion coefficient profiles, could readily be integrated to obtain permeability coefficients, via the inhomogenous solubility-diffusion (ISD) model [33] (see eq. 7 and methods).

We calculated the local diffusion coefficient along the permeation coordinate from the autocorrelation function of the positions from the same biased, i.e. umbrella sampling, MD simulations which we also used to compute the free energy profiles of permeation [34] (Figure 4A). As expected, for all molecules, the local diffusion coefficient dropped by approximately one order of magnitude at the more viscous interior of the membrane compared to the bulk water. The reduction in local diffusivity was already observed before the molecules entered the membrane, possibly due to solvent-mediated interactions between the molecules and the membrane [35]. The difference in the local diffusion coefficient between the compounds was less pronounced than the changes in free energy. Diffusion coefficients were found to be inversely correlated with the size of the molecules (Figure 4B,C). Interestingly, for small compounds such as AEMA and NIPAM, the local diffusion coefficient rose again in the center of the membrane where the acyl chains are less densely packed and friction is therefore reduced (Figure 4A). This was not the case for the larger molecule RHODB-SL that still interacted with the densely packed regions of the membrane. Although the observation that diffusion reduces with viscosity and molecule size is to be expected, the simulations could also spatially resolve the diffusive behavior of the resist inside the lipid bilayer, something that is simply not accessible in experiments. This once again underlines the high resolution that molecular dynamics provides in the investigation of membrane permeation by small molecules. More importantly, the diffusion coefficient calculations enabled us to establish the connection to permeation kinetics as explained as follows.

**Figure 4:**
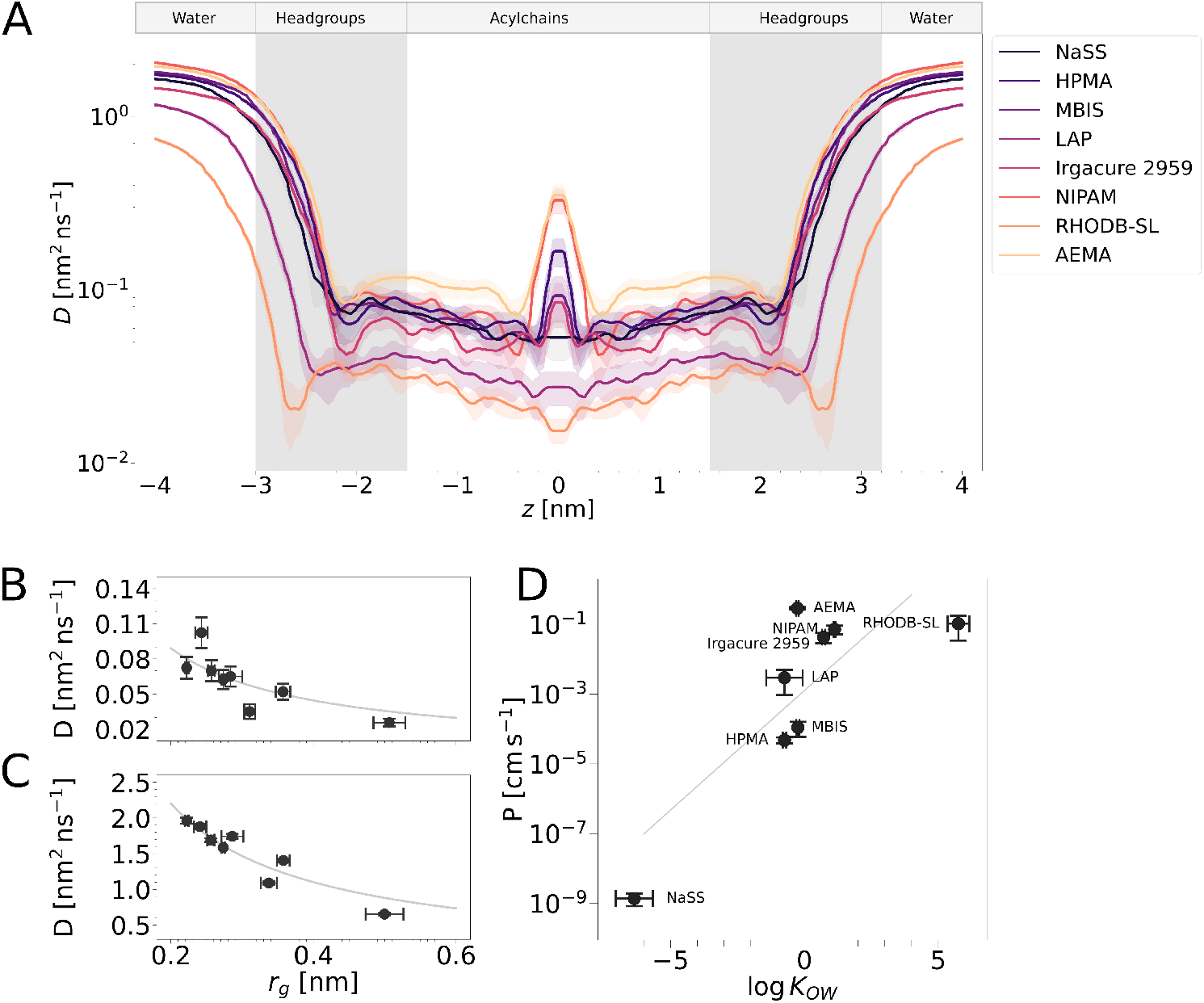
Kinetics of permeation of photoresists across a POPC lipid bilayer. A) Diffusion coefficient *D* as a function of the *z* position of the photoresists. Lines indicate the smoothed result and the fill the standard error from three replicates (see methods for details). The approximate location of bulk water, headgroups and hydrophobic acyl chain regions is indicated. B–C) *D* is presented as a function of the radius of gyration *r*_*g*_ of the photoresists in the membrane (*z* = 1 nm, B) and the bulk water (*z* = 3.8 nm, C). Time-average± standard deviation is shown for *r*_*g*_. The lines display fits of the form *D* = *k/r*_*g*_, with resulting parameters *k*_*z*=1.0 nm_ = 0.018 ± 0.002 nm^3^ ns^−1^ and *k*_*z*=3.8 nm_ = 0.440 ± 0.016 nm^3^ ns^−1^. D) Correlation (r=0.82) between computed permeabilities *P* and log *K*_*OW*_, along with their uncertainties, is shown (see methods for details on the statistical uncertainty estimation).

The free energy and the diffusion profiles were joined in the ISD model to obtain permeation coefficients. In line with Overton’s rule [36], we found that the permeability of the photoresists correlate reasonably well with their hydrophobicity as quantified by their log *K*_*OW*_ value (*r* = 0.81 over all resists in Figure 4D). Thus, the simple notion that more hydrophilic molecules will less efficiently permeate the membrane appears to be a reasonable criterion to judge cytocompatibility of the photoresist used for 3D bioprinting. The observed relation between log *K*_*OW*_ and permeability strongly resembles the one previously found for small solutes [19, 37], further stressing the robustness of our computational protocol. As previously suggested, the estimated permeability of the photoresist NaSS is more than four orders of magnitude lower than for the other photoresists, supporting the idea that this molecule would not permeate the lipid membrane on experimental timescales. HPMA and MBIS follow with permeabilities that are at least two orders of magnitude smaller than that of the next more permeable molecule LAP. Finally, AEMA, NIPAM, Irgacure 2959, RHODB-SL and LAP could be classified into the group of molecules with highest permeability.

Experimentally, considering the printing success rate over time as a proxy for the intravesicular concentration of the initiator allowed the determination of the permeability for LAP [11]. In this study, we assessed the permeation rate of RHODB, by directly using its spectroscopic properties to measure its influx in the GUV (Table 2). Due to the velocity in which the experimental system saturated (Figure 5B), for pure RHODB a lower boundary could only be estimated. In our previous work, we determined the permeation kinetics of RHODB together with LAP [11]. In the presence of LAP, RHODB exhibited an about one order of magnitude slower influx into the GUVs (Table 2).

**Table 2:** Permeability for selected photoresists. Comparison of permeation rates obtained from experiments and biased molecular dynamics simulations.

| Name | Permeability (MD)<br>(cm/s) | Permeability (Experiment)<br>(cm/s) |
| --- | --- | --- |
| RHODB-SL | $(104 \pm 71) \times 10^{-3}$ | $\geq 10^{-5}$ |
| RHODB-SL (with LAP): | NA | $(2.301 \pm 0.016) \times 10^{-7}$ (From ref. [11]) |
| LAP | $(3.0 \pm 2.1) \times 10^{-3}$ | $(0.987 \pm 0.325) \times 10^{-7}$ (From ref. [11]) |

**Figure 5:**
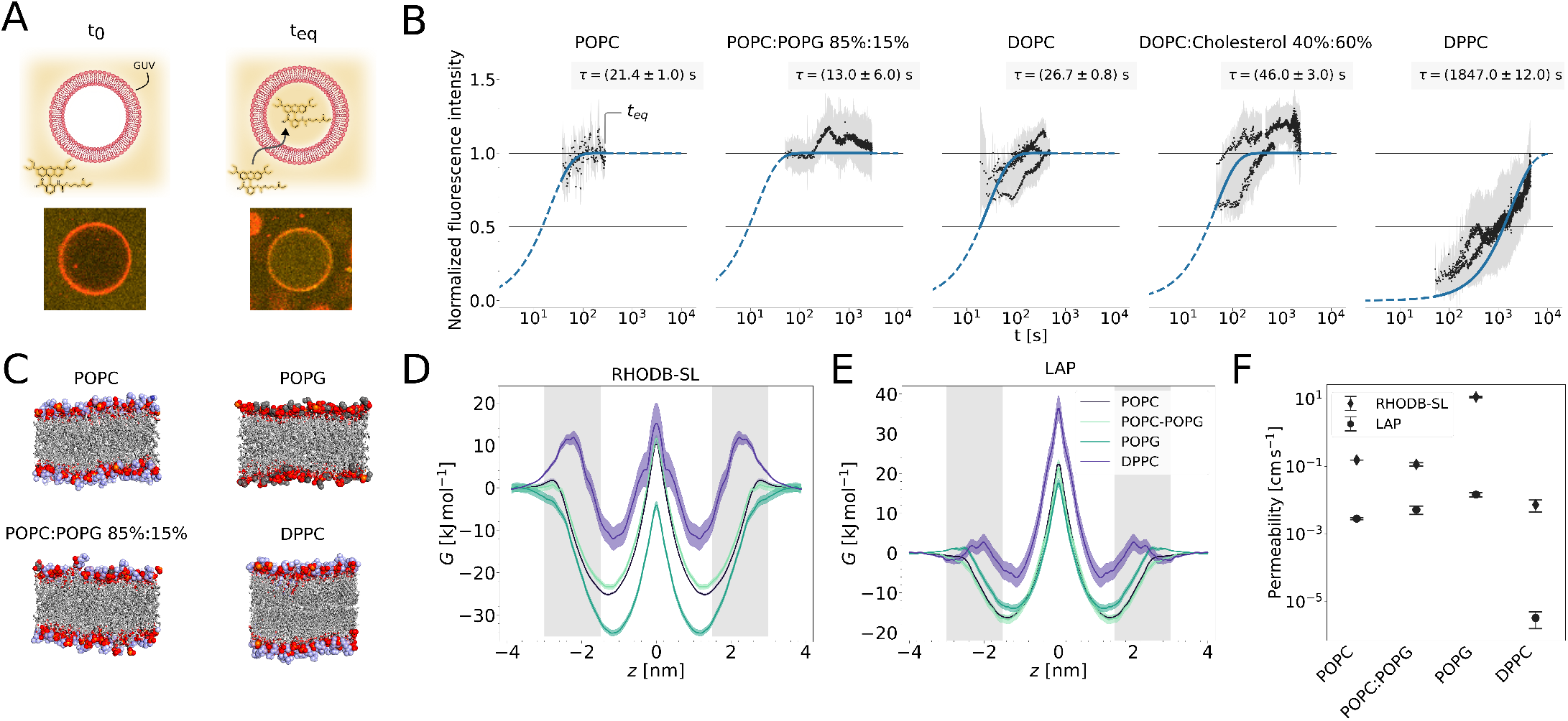
Influence of membrane composition on photoresist permeation. A) Top: Experimental principle of the influx experiments with the fluorescent photoresist Rhodamine B acrylate (RHODB) into giant unilamellar lipid vesicles (GUVs, red) of varying lipid composition. The fluorescence within the GUVs was measured over time as surrogate for the intra-vesicular concentration of the dye until an equilibrium was reached, i.e. when no change in fluorescence in the inside was observed. Bottom: Representative images of a POPC (85 %) POPG (15 %) GUV at the start of the measurement (*t*_0_) and when equilibrium was reached (*t*_*eq*_). B) Results of influx experiments with Rhodamine B for different membrane compositions (POPC N=2, POPC-POPG N=1, DOPC N=4, DOPC-Cholesterol N=2, DPPC=2). For every time point *t* we report the mean fluorescence intensity of all *n* GUVs that were in the field of view (black dots, 2 *< n <* 52) normalized to an averaged fluorescence intensity outside of all GUVs and the standard deviation of the measurement (gray shaded area). The normalized fluorescence intensity 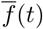 is proportional to the intra-vesicular concentration of the dye. Non-linear least squares fits of the form 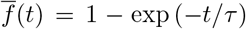 are shown as blue solid lines in the time period that was used for the fit and as dashed lines in the rest of the time period. Fit values *τ* are indicated in the plots. C) Snapshots of membranes considered in this study are shown. They consisted of neutral 1-Palmitoyl-2-oleoylphosphatidylcholine (POPC), negatively charged 1-Palmitoyl-2-oleoylphosphatidylglycerol (POPG), or fully saturated 1,2-Dipalmitoylphosphatidylcholine (DPPC) lipids. Acyl chains are depicted as gray sticks and headgroups as spheres (red-oxygen, orange-phophorous, blue-choline headgroup, gray-glycerol carbons). In the simulations the bilayers were fully solvated by explicit water molecules and ions (not shown for clarity). D–E) Free energy profiles *G*(*z*) for the fluorescent dye RHODB-SL (D) and the photoinitiator LAP (E) at the indicated lipid bilayers. Same format was used as in Figure 3. F) Computed permeabilities for the molecules RHODB-SL (black squares) and LAP (black circles) in dependence on the membrane composition.

Both in the computed and the experimental permeabilities, we observed this trend, i.e. LAP permeates at a two orders of magnitude slower rate than RHODB-SL (Table 2). However, the computational values overestimate the experimentally observed permeabilities by several orders of magnitude. Several computational and experimental factors might contribute to these deviations. A significant overestimation in the permeability predicted by the ISD model, compared to the experimental one, was also observed in other studies of small solutes [13, 19]. This is likely due to the underestimation of the viscosity of the TIP3P water model [38] and hydrocarbons [39], leading to intrinsically faster dynamics in the simulations. Moreover, as mentioned above, only an upper boundary of the permeation rate could be determined for pure RHODB, in combination with an estimate which was one to two orders of magnitude lower than this boundary, but conditioned to the presence of LAP in the solution (Table 2). This suggests that interactions between these two molecules influence the permeation, a feature that is not taken into account in the ISD model. More generally, any interaction between solutes is not accounted for in this simplified representation, a factor which likely contributes to the deviation between simulations and experiment.

However, despite its limitations in the quantitative prediction of permeation rates, ordering the photoresists qualitatively by their permeabilities still provides valuable information. Previously, this has been used as a bioavailability criterion to classify drug-like molecules [16]. Similarly, we propose to utilize the relative change in permeability to assess the likelihood of lipid membrane permeation on experimentally relevant time scales, i.e. during a bioprinting experiment. Membrane permeation may be the root cause for cytotoxicity or alter the cells in other ways, which is why awareness about this effect is crucial for the bioprinting community. For example, Xu et al. [40] report that Irgacure 2959 exhibits a higher cytotoxicity than LAP when used as photoinitiator in laser assisted 3D bioprinting. This correlates with our predicted membrane permeabilities: Irgacure 2959 permeates faster than LAP (Figure 4D). We therefore give a possible explanation for the observed level of cytotoxicity of these two molecules in terms of their ability to cross the lipid bilayer.

### 2.5 The influence of membrane composition on resist energetics and permeation

Finally, we investigated the effect of lipid composition on the energetics and permeation of the photoresists. This is of particular interest since biological membranes exhibit a complex lipid composition, featuring a rich variety of lipid headgroups, including charged groups, and acyl chains of distinct lengths and saturation. It is expected that these factors influence the permeability of solutes [19]. Here, we focused on three key bilayer properties, namely the electrostatic charge of the headgroup, the presence of sterol molecules, and the saturation of the acyl chains.

First, we experimentally assessed the permeability of the fluorescent dye RHODB into GUVs of varying lipid composition by means of influx experiments into GUVs observed by confocal microscopy (Figure 5A). We considered the fluorescence of RHODB inside the GUV as a proxy of the intra-vesicular resist concentration. Accordingly, monitoring the kinetics of the fluorescence influx indicated how fast this molecule crossed the membrane along the concentration gradient. As a reference, we considered GUVs composed of POPC lipids.

To study the effect of charge, we considered a mixture of POPC with the negatively charged lipid 1Palmitoyl-2-oleoyl-phosphatidylglycerol (POPG) (85:15 POPC:POPG mol ratio) resembling the typical charge distribution of the lipids in the inner leaflet of cell membranes [41, 42]. Note that the POPG concentration in lipid bilayers is typically about 1–2 % and in monolayers, such as in the lung surfactant, as high as 9 % [43]. However, POPG was used here as a rather generic charged phospholipid, allowing to study the effect of the charge of the headgroups *in silico* and *in vitro*. In the experiments, using this amount of POPG lipids did not significantly vary the permeation rate (Figure 5B), highlighting that the hydrophobic barrier, imposed by the acyl chains, rather than the charge of the head groups, dominates the permeation kinetics. Furthermore, we investigated the influence of introducing cholesterol into a 16:0 PC 1,2-dipalmitoyl-sn-glycero-3-phosphocholine (DOPC) GUV (i.e ∼ 40:60 DOPC:Cholesterol mol ratio). Note that this corresponds to a high concentration compared to what is typically found in plasma membranes [44], although still manifested in some instances, such as the eye lens [45, 46, 47]. Despite the high cholesterol content, we observed only a slight decrease in permeation kinetics (Figure 5B). In addition, to explore the effect of the lipid ordering imposed by the acyl chain saturation, we considered GUVs composed of fully saturated 1,2-dipalmitoyl-sn-glycero-3-phosphocholine (DPPC) lipids. This composition significantly slowed down the permeation compared to the other GUV compositions analyzed here (Figure 5).

In a next step, we analyzed the effect of lipid composition in MD simulations, by considering pure POPC, POPG, and DPPC bilayers as well as a binary mixture (85:15 mol ratio) of POPC and POPG (Figure 5C). We focused on the experimentally version of the photoresist, RHODB-SL, in order to be able to directly compare simulations with experiments (Figure 5D,F). We also investigated the behavior of the ionic photoinitiator LAP to discern the effect of the charged headgroups on its energetics and permeation (Figure 5E,F). As already observed in the experiments, introducing 15 % mol of the charged lipid POPG did not significantly vary neither the energy profile (Figure 5D) nor the permeability derived from it for RHODB-SL (Figure 5F). Similarly, LAP remained practically insensitive to this change in lipid composition (Figure 5E,F). As described, we observed an important electrostatic interaction of the positively charged lithium ion, bridging the ternary interaction between the photointiator LAP, positively charged ions and the negatively-charged phosphate groups (Figure 3F,G). Replacement of the positively-charged choline by the neutral glycerol moiety (in 15 % of the lipids) did not seem to alter this interaction, at least not to an extent that would influence the energetics and the permeation. An increase to 100 % POPG concentration was necessary to induce a noticeable effect (Figure 5D,F). We predict slightly higher permeabilities for both LAP and RHODB-SL for the pure POPG bilayer compared to the reference POPC bilayer. Presumably, this is due to the less dense packing, i.e. higher area per lipid, imposed by the electrostatic repulsion of the phosphatidylglycerol (PG) headgroups. While this highlights the importance of lipid packing on the membrane permeability of the photoresists, a pure POPG membrane represents an extreme scenario that could not be attained experimentally and that is far from the membrane composition of cells. In contrast to POPG, for a fully saturated DPPC lipid bilayer in the gel phase at 300 K, the calculated permeability of both resists RHODB-SL and LAP was significantly reduced (Figure 5F). This was a consequence of the higher energy barrier in the middle of the membrane and a lowering of the energy well at the acyl chain-headgroup interface (Figure 5D,E). In the gel phase, the DPPC lipids displayed a pronounced ordering of the acyl chains leading to a more dense packing of the lipids compared to POPC (see order parameter and area per lipid in Figure S4 and Table S3). We thus attribute these changes in energetics and kinetics to the difficulty of accommodating the resist molecules in the tightly packed gel environment provided by the DPPC lipids.

Both our experiments and simulations consistently show that augmenting the acyl chain saturation, while turning the bilayer into a gel, significantly reduced the permeation of RHODB. Our simulations extended this finding to a second resist component, namely LAP, which cannot be tracked directly in experiments. Thus, low permeability across gel-like membranes is suggested to be a general feature that applies not only to small neutral solutes [19] but also to photoresist molecules. Concerning sterols, a more moderate reduction in the permeability was observed for cholesterol-containing membranes consistent with previous reports [19]. Our combined approach shows that the introduction of charged lipids in a physiologically relevant amount (15 %) did not induce an appreciable effect on the permeability of RHODB or the ionic LAP. Altogether, our results demonstrate key features of membrane composition to specifically impact the permeation of photoresist molecules.

## 3 Conclusions

Here, we have used MD simulations, together with octanol-water partitioning experiments and influx assays in GUVs to study the interplay between common photoresists used for bioprinting and biological lipid membranes. By this combined approach we quantitatively classify a representative set of photoresist molecules according to their likelihood to permeate through or to reside in biological lipid bilayers. A central aspect of 3D bioprinting is the cytocompatibility of the used molecules. In this respect, the observed free energy barriers reveal the extent by which the hydrophobic core of the membrane imposes a barrier to the photoresist molecules and prevent them from crossing the membrane and therefore entering the cytosol. In addition, these data showed the existence of energetically-favorable accumulation sites directly at the acyl chain-headgroup interface. By combining diffusion coefficients and free energy profiles, a broad range of permeation rates for the studied compounds, spanning almost eight orders of magnitude, was observed. Furthermore, permeabilities were found to correlate well with the determined octanol-water partition coefficients, i.e. they followed Overton’s rule. A major result of our study is that many photoresist components used in 3D bioprinting, such as the monomers (AEMA, NIPAM), the photoinitators (LAP, Irgacure 2959), and the fluorescent dye (RHODB-SL) are typically membrane-permeable on the timescale of a 3D printing experiment. The photoresists NIPAM, LAP, and RHODB-SL further tend to accumulate at the headgroup region of the membranes. This might impact cellular behavior or even induce cytotoxicity. Besides, we predict other photoresist molecules, such as NaSS to be largely membraneimpermeable. Using membrane impermeable photoresists might reduce the impact of 3D bioprinting on cellular homeostasis. Thermodynamic and kinetic characterization can therefore guide the choice of photoresists for 3D bioprinting experiments.

The reduced permeability across highly-order gel-like lipid bilayers, previously observed for small neutral solutes, was also found here to apply to photoresist molecules. This provides an initial glimpse into how lipid composition influences the permeation behavior of photoresists and serves as a basis for future studies of membranes of increasing complexity.

Overall, our data provides the molecular basis of the interaction of photoresist molecules with model biological lipid bilayers. Resists that enter the cells on experimentally relevant time scales are expected to interfere with cellular processes. In particular, the generation of free radicals inside the cell during printing is cytotoxic. Therefore, we recommend the use of non-cell-permeable resist formulations in bioprinting experiments. Accordingly, our computational approach in combination with experimental data, by quantitatively predicting their energetics and permeability, will help to rationally select suitable bioinks for different 3D bioprinting applications.

## 4 Methods

### 4.1 Parametrization of photoresist molecules for MD simulations

The parameters for the photoresist molecules were generated with the ligand reader and modeler tool on the CHARMM-GUI website (https://www.charmm-gui.org/, Version 3.7) [48, 49] using the CHARMM36 general forcefield (CGenFF, version 2.5.1) [50]. SMILES codes of these molecules (Table S1) were considered as input for this purpose.

The confidence of the obtained parametrization was assessed with the heuristic penalty score provided in the CHARMM-GUI output (Figure S2A). For LAP and RHODB-SL we observed several high (*>* 50) scores, associated with less confidence in the assignment of the respective CGenFF parameters, specifically in proximity to their phosphinate and spiro moieties, respectively, for which no exactly equivalent atom types exist in the CGenFF. For these two molecules optimized structures, charges, and selected bonded parameters were also assessed by means of density functional theory (DFT) calculations.

#### DFT calculations

For the molecules LAP and RHODB-SL, with DFT calculations, we verified their structures, charges, and selected dihedrals, for which the CHARMM-GUI penalty exceeded 50 (Figure S2A). The computations were set up using the FFParam package [51] and carried out with Gaussian 09 [52]. To optimize the structures of RHODB-SL and LAP, we performed calculations at the Møller-Plesset second order theory [53] (MP2) with the 6-31G* basis set [54]. We further tested whether assigning these point charges improved the correspondence between log *K*_*OW*_ values from experiments/RDKit and MD simulations (LAPQM/RHOSLQM, Figure S2B,C). This was generally not the case. Similarly, the minimal adjustment of few selected charges that displayed the largest variation between the CHARMM-GUI and the QM data sets (LAPQMSEL, RHOSLQMSEL) or stiffening of selected dihedrals (LAPSTIFF, RHOSLSTIFF) did not improve the agreement between simulations and experiments. Overall, we found the obtained parameters to be in good agreement with the original output from CHARMM-GUI.

#### Equilibration of photoresist molecules

LAP and NaSS coordinate a lithium and a sodium ion, respectively. We manually positioned the ion near the coordination site (Figure 1). Unless stated otherwise, we always included such a coordinating ion for each LAP or NaSS molecule. We minimized the energy of the parameterized resists (and coordinated ion in the case of LAP and NaSS) in vacuum. We next solvated the resists in a cubic box filled with water molecules of dimensions of 5 nm × 5 nm × 5 nm and again, we performed energy minimization. Afterwards, we thermalized the system in 500 ps to a temperature of 300 K, followed by a 500 ps pressure equilibration with a reference pressure of 1 bar. Finally, we carried out MD simulations of the solvated resists for 5 ns at 300 K and 1 bar. After equilibration, we extracted the coordinates of the resist molecules for the following steps.

### 4.2 Simulation parameters and algorithms

All-atom molecular dynamics (MD) simulations were performed using GROMACS (version 2021) [55]. The CHARMM36 force field was used for the lipids [56] and octanol [57], the CGenFF for the resists (see section 4.1 above), the CHARMM-TIP3P model [58, 59] for the water molecules, and the default CHARMM parameters for the ions [60]. We minimized the energy using the steepest descent algorithm. Bonds involving hydrogen atoms were constrained with the LINCS algorithm [61], while both bonds and angles involving hydrogens of water molecules were constrained with SETTLE [62]. We employed the Verlet-buffer neighbor list scheme [63] with a cut-off of 1.0 nm and a van der Waals force switch up to 1.2 nm. Temperature was kept constant with the Nosé-Hoover temperature coupling scheme [64, 65] during production runs and the Berendsen one [66] during equilibration or relaxation steps (relaxation time 𝒯_*T*_ = 1.0 ps). Membrane and solvent (i.e. resists, water molecules and ions) were coupled separately. Pressure was maintained constant employing the Parinello-Rahman barostat [67, 68] (Berendsen during equilibration and relaxation) with a pressure relaxation time 𝒯_*T*_ = 5.0 ps and compressibility of 4.5 • 10^−5^ bar^−1^. To treat long-range electrostatic interactions we employed the Particle-Mesh-Ewald scheme [69, 70].

### 4.3 Computational prediction of partition coefficients

#### Partition coefficients from non-equilibrium MD-based free energy calculations

The partition coefficient quantifies the distribution of resist molecules in biphasic octanol-water systems and is proportional to the free energy of transferring each resist from octanol (*o*) to water (*w*), Δ*G*_*o*→*w*_ (Figure 2A). Non-equilibrium free energy calculations were performed to estimate this transfer free energy by using the thermodynamic cycle shown in Figure 2A. By calculating the solvation free energy for each resist in octanol, Δ*G*_∅→*o*_, and in water, Δ*G*_∅→*w*_, separately, we obtained Δ*G*_*o*→*w*_ = Δ*G*_∅→*w*_ − Δ*G*_∅→*o*_. Accordingly, the partition coefficient reads:

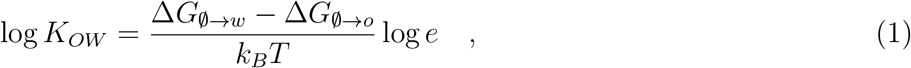

with *k*_*B*_ the Boltzmann constant, *T* the temperature, and *e* the Euler number. The solvation free energies in each solvent, *s* = (*o, w*), were estimated through thermodynamic integration [71]. The dissolved (in vacuum) and solvated states were coupled by a *λ* coordinate, such that solute-solvent interactions were switched on for *λ* = 0 and switched off for *λ* = 1 (Figure 2A). Accordingly, transitions *λ* = 1 → 0 resembled the solvation process. For practical purposes, we estimated the free energy associated to the transition *λ* = 0 → 1, corresponding to the desolvation process (Δ*G* ≡ −Δ*G*_∅→*s*_). By integrating the derivative of the Hamiltonian *H* with respect to the coordinate *λ*, the non-equilibrium work associated to the solvation transition was obtained as 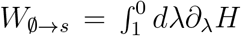 and the work related to the desolvation transition as 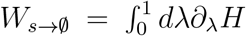. Free energies were computed from forward *P*_*s*→∅_(*W*) and backward *P*_∅→*s*_(*W*) work distributions using the Crooks fluctuation theorem [72]:

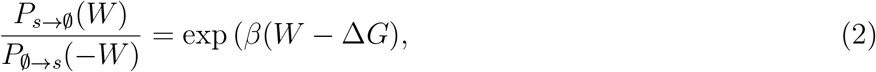

where *β* = 1*/*(*k*_*B*_*T*). Starting from both work distributions, Bennett’s acceptance ratio was used as a maximum likelihood estimator to numerically estimate the change in free energy Δ*G* upon solvation, as proposed by Shirts [73]:

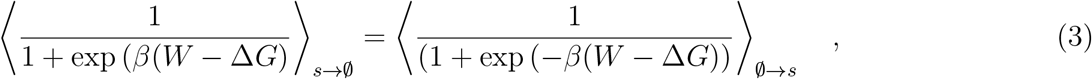

where ⟨…⟩ denotes the average taken over an ensemble of simulations, further allowing the quantification of uncertainty through bootstrapping.

We solvated each photoresist molecule by either 300 octanol or 4000 water molecules. In case of charged photoresist molecules, such as LAP or NaSS, one extra ion was added to maintain the neutral net charge of the system. Before performing thermodynamic integration, we simulated the solvated photoresist under equilibrium conditions during 50 ns, in both states, *λ* = 1 and *λ* = 0, having the Coulomb and Lennard-Jones interactions between resist and solvent turned on or off, respectively. Next, we extracted 200 conformations (equally spaced by 200 ps) during the last 40 ns of each equilibrium simulation as starting configurations for the non-equilibrium transitions. Accordingly, 200 forward (*λ* = 0 → 1) and 200 backwards (*λ* = 1 → 0) transitions were simulated for 1 ns each, using the implementation of free energy calculation suite of GROMACS. A Beuter soft-core potential was applied to prevent singularities when jointly switching Lennard-Jones and Coulomb interactions [74] (*α* = 0.3, *σ* = 0.25 nm, and a soft-core power of 1 parameters were used). Free energy of solvation, together with its uncertainty by means of bootstrapping, was computed for each resist in both solvents using the implementation of the *pmx* package [75, 76]. log *K*_*OW*_ was subsequently obtained using eq. (1) and its uncertainty by propagating the error of the Δ*G* bootstrap estimates.

#### Partition coefficients from atomic contribution approach

For the non-ionic molecules, their octanol-water partition coefficients were also predicted from the respective Simplified Molecular Input Line Entry Specification (SMILES) codes with the atom-typing based methodology [25] implemented in the python package RDKit (v. 2022.3.2) [24].

### 4.4 Free energies, diffusion coefficients, permeabilities, and ion-molecule binding frequencies from Umbrella sampling MD simulations

#### Lipid bilayer simulation setup

The membrane lipid bilayer systems were generated using the membrane builder input generator on the CHARMM-GUI website [48, 77, 78]. The membranes were composed of a total of 162 lipids with a symmetric lipid composition between leaflets (Table S2). All membranes were hydrated with 70 TIP3 water molecules per lipid and neutralized with 0.15 mM NaCl. For systems containing PG charged lipids, additional sodium ions were added to neutralize the system. The membranes were first equilibrated with the default workflow provided by CHARMM-GUI, starting with an energy minimization step followed a sequence of equilibration steps gradually removing position and dihedral restraints on the lipid head groups. Subsequently, we performed a 1 *µ*s production run with a step size of 2 fs under a NPT ensemble, by keeping the temperature constant at 310 K the pressure at 1 bar. Snapshots of the membranes at 800 ns, 900 ns and 1000 ns were taken as initial configurations for the umbrella sampling simulations.

#### Umbrella sampling MD simulations

To obtain free energy profiles of membrane permeation of the photoresists, we conducted umbrella sampling MD simulations. A harmonic umbrella potential of 1000 kJ mol^−1^ nm^−2^ was applied on the center of mass of each solute along the permeation coordinate, i.e. perpendicular to the membrane surface (named *z* coordinate). We considered multiple umbrella simulations spaced by 0.025 nm while omitting a 1.2 nm buffer region close to the edge of the box to reduce boundary effects. Aiming to reduce the computational costs of the sampling procedure, we followed the approach proposed by Nitschke and coworkers [32] and sampled multiple umbrella windows per simulation. To this end we placed several photoresists in the same simulation box. Nitschke et al. showed that the placement of several solutes in one system does not significantly affect the accuracy of the computed free energy profile provided that they are sufficiently separated from each other [32]. Accordingly, we set the resist-resist distance Δ*Z*, defined by the minimal distance between any part of the two molecules plus an extra buffer distance Δ*Z*_*B*_ = 2.0 nm, to Δ*Z* ≈ 2.5 nm (RHODB-SL: Δ*Z* ≈ 3.5 nm). To maximize the number of sampled windows per simulation, the molecules were placed in two separate columns with an offset of Δ*Z*_*B*_*/*2 between the columns along the permeation (*z*) coordinate and a lateral distance of half of the box width between the columns along the *y* coordinate (see Figure 1). In order to prevent interactions of solutes from different columns, each column of resist molecules was maintained in its own half of the box. A flat bottom harmonic potential applied on each resist molecule along y coordinate parallel to the membrane surface was used for this purpose (flat bottom distance of 1.0 nm and elastic constant of 2000 kJ mol^−1^ nm^−2^). We placed the molecules in the center of the respective umbrella potentials using the most central atom of the membrane system as reference point. After inserting the solutes in the membrane system, the systems were equilibrated while constraining the positions of the molecules as explained in the following.

First, we removed the overlap between resists and lipids and water by gradually switching on Lennard-Jones and electrostatic parameters of the resists in a Langevin stochastic dynamics simulation of 10 ps (at 310 K and with a friction coefficient for each atom given by its mass divided by 1.0 ps). The GROMACS free energy code was used for this purpose, including soft core parameters set to *α* = 0.3 and *σ* = 0.25 nm to avoid singularities. To prevent the rings from being penetrated by lipid tails, we added a dummy atom to the geometric center of all aromatic rings of Irgacure 2959, NaSS, LAP and RHODB-SL. The dummy had Lennard-Jones parameters corresponding to an aromatic carbon atom and neutral charge. It was also bound to the carbon atoms in the respective ring. Subsequently, the system was energy minimized, followed by a 25 ps NVT thermalization at 310 K and 2 ns NPT equilibration at 1 bar. During all these steps, position restraints were applied on the atoms of the resist (elastic constant of 1000 kJ mol^−1^ nm^−2^) and on the phosphorous atoms of the lipids (elastic constant of 100 kJ mol^−1^ nm^−2^), and dihedral restraints at the headgroup tail interface of the lipids (elastic constant of 2.5 kJ mol^−1^). At this point, dummy atoms were removed and position and dihedral restraints lifted, to subsequently run 50 ns (RHODB-SL: 100 ns) long umbrella simulations for every window. We saved coordinates of the trajectories every 100 ps. This procedure was carried out three times, starting with different initial configurations of the lipid bilayers. The overall cumulative sampling, considering all resists and lipid compositions, was more than 72 *µ*s.

The free energy profile was determined from the umbrella sampling simulations, via the GROMACS tool gmx wham [79], by employing the weighted histogram analysis method (WHAM) [80]. The reported free energy estimates represent the estimates when considering the aggregated information from all replicates (POPC: 3 replicates; POPC-POPG, POPG, DPPC: 1 replicate), where —if applicable— the replicates represent simulations starting from different initial configurations (see above). Statistical uncertainty of the free energy profiles was quantified by two hundred fold Bayesian bootstrapping runs of the combined histograms from all replica as implemented in the gmx wham tool [79]. To facilitate the comparison, all free energy profiles are reported relative to the reference in bulk water (3.5 nm − 4.0 nm). Convergence in the free energy profiles was assessed by backwards block averaging. Accordingly, the first 20 ns (RHODB-SL: 50 ns) of each umbrella window was discarded from the WHAM calculation (Figure S5,S6).

#### Diffusion coefficient

The diffusion coefficient of a molecule in umbrella sampling simulations can be determined with the simplified version of the Wolf-Roux [81] estimator proposed by Hummer [34], which has been employed for this purpose in previous studies [14]. In short, for each umbrella we computed the diffusion coefficient as

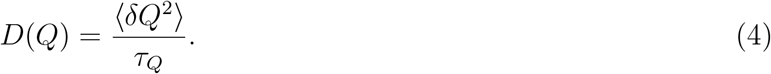

Here, *Q* is the average biased position over time of the resist molecule along the permeation coordinate. ⟨*δQ*^2^⟩ is the variance and *τ*_*Q*_ the correlation time of *Q. τ*_*q*_ is defined as the integral over time of the autocorrelation function ⟨*δQ*(*t*)*δQ*(0)⟩ by the variance [34]:

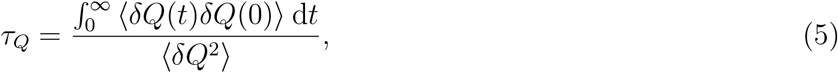

with *δQ*(*t*) being the deviation of the molecule’s position along the permeation coordinate from its average position:

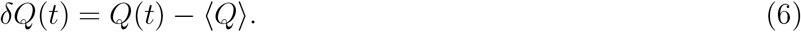

In practice, we evaluated the integral up to the first time point at which the autocorrelation function reached zero. The coordinates *Q* were extracted from the simulations for each umbrella. The autocorrelation function was computed with the gmx analyze GROMACS tool. Diffusion coefficients from all windows yielded a diffusion coefficient profile along the permeation coordinate: *D*(*z*), where *z* is the average position of the resist along the permeation coordinate. Diffusion coefficient profiles were smoothed by applying a Gaussian filter with a width of five standard deviations (Figure S7,S8). The obtained local diffusivities were linearly interpolated to the same permeation coordinates as the free energy profile. The reported diffusivity profiles correspond to the average from three smoothened replicates. Errors of the local diffusivity were estimated by running the same analysis separately on five subsamples of equal length of the trajectories. The standard deviation of the obtained local diffusion coefficients is the error for one replicate. If multiple replicates were performed for one condition (i.e. the POPC membrane), we report the estimated error from the propagation of uncertainties of the individual replicates.

#### Permeability coefficient

We employed the inhomogenous solubility-diffusion (ISD) model [33] to derive the permeability coefficient *P* from the previously calculated free energy and diffusion coefficient profiles:

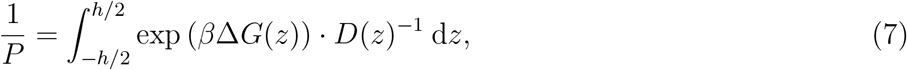

where *h* denotes the thickness of the membrane (POPC, POPG, POPC-POPG: 6.4 nm based on the z coordinate where the density of the ammonium functionality of the POPC membrane’s headgroups reaches zero, 7.6 nm for DPPC), *D*(*z*) the diffusion coefficient, Δ*G*(*z*) = *G*(*z*) − *G*_*w*_ the position-dependent free energy with respect to the bulk water and the thermodynamic *β* = 1*/*(*k*_*B*_*T*). Note that *z* = 0 corresponds to the center of the membrane. We numerically evaluated the integral in eq. 7 using the trapezoid rule. The reported error *e*(*P*) was obtained by propagation of uncertainties of the individual points of the free energy profile and diffusion profile (Supplementary text S3).

#### Ion-molecule binding probability

For the charged photoresists LAP and NaSS we examined their position dependent interaction probability with positive ions. For every molecule and every frame of the trajectory, we determined the position *z* of the molecule relative to the center of membrane and the minimal ion-molecule distance *d*, separately for every ion-species (LAP: Lithium, Sodium, and Chloride, and NaSS: Sodium and Chloride). If the distance was below *d <* 0.5 nm, the frame was counted as an binding event. For every binned *z* position, we report the probability of observing an ion bound to the molecule, *P*_bound_, as the number of frames in which we observed a binding event divided by the total number of frames of the simulation. The bin width was Δ*z* = 0.25 nm.

### 4.5 Materials for experiments

N-Isopropylacrylamide (NIPAM), N,N’-Methylenebis(acrylamide) (MBIS), Lithium phenyl-2,4,6-trimethylbenzoylphosphinate (LAP), 2-Hydroxy-4’-(2-hydroxyethoxy)-2-methylpropiophenone (Irgacure 2959), sucrose, glucose, poly(vinyl alcohol) (PVA, M_w_ 13 000 to 23 000) were purchased from Sigma-Aldrich. N-(2Hydroxypropyl)methacrylamide (HPMA), Acryloxyethyl thiocarbamoyl Rhodamine B (RHODB), Cholesterol were purchased from PolyScience. 1-octanol and Corning UV 96 well plates were purchased from Merck Millipore. 1,2-dioleoyl-sn-glycero-3-phosphocholine (DOPC), 1,2-dipalmitoyl-sn-glycero-3-phosphocholine (DPPC), 1-palmitoyl-2-oleoyl-sn-glycero-3-phospho-(1’-rac-glycerol) (sodium salt) (POPG) and 1-palmitoyl-2-oleoyl-glycero-3-phosphocholine (POPC) were purchased from Avanti Polar Lipids. Atto633-1,2-Dioleoyl-sn-glycero-3-phosphoethanolamine (Atto633-DOPE) was purchased from ATTOTEC GmbH. Buffer solution pH 13 was purchased from Carl Roth. Indium tin oxide (ITO) coated glass coverslips were purchased from Visiontek Systems Ltd. Poly(dimethylsiloxane) (PDMS, Sylgard 184 from DOW Corning) and petri dishes (Falcon 100 mm cell culture dish, Corning) were purchased from VWR. A 3 mm biopsy puncher (Harris Uni-Core, 3mm) was purchased from Electron Microscopy Sciences.

### 4.6 Octanol-water partition coefficient determination via osmolality measurements

For HPMA, NIPAM and MBIS, the octanol-water partition coefficient was determined using an osmometer. Briefly, HPMA, NIPAM and MBIS were dissolved in Milli-Q water at 43.9 g l^−1^, 41.7 g l^−1^ and 30.1 g l^−1^, respectively. 15 *µ*l of each solution were separated for osmometer measurements. Another 200 *µ*l of each solution were mixed with additional 200 *µ*l of 1-octanol, vortexed for 2 min and centrifuged for 1 min at 2000 g using a table top centrifuge (Corning Mini Centrifuge, Corning). 15 *µ*l of the aqueous solutions were extracted from the phase separated liquids for osmolality measurements. To obtain the partitioning coefficient, the obtained osmolality was first substracted from the initial osmolality and then divided by the initial osmolality of the respective solution prior to the octanol addition. All osmolality measurements were conducted using an Osmomat 030G (Gonotec GmbH). Prior to usage, the osmometer was calibrated using calibration solutions of 0 mOsm kg^−1^ and 300 mOsm kg^−1^ (Gonotec GmbH).

### 4.7 Octanol-water partition coefficient determination via absorption

For LAP, Irgacure 2959 and RHODB, the octanol-water partition coefficient was determined via UV-vis absorption measurements. Briefly, LAP was dissolved at 10, 7.5, 5 and 2.5 g l^−1^ in Milli-Q water. Irgacure 2959 was dissolved in Milli-Q water at 0.2 g l^−1^ and diluted five times in a serial dilution by a factor of two each time. RHODB was dissolved in DPBS or pH 13 buffer solution at 1 g l^−1^ and diluted 11 times in a serial dilution by a factor of two each time. Using UV-96 well plates, the absorbance spectra for all concentrations including the dilution steps were recorded from 200 or 300 nm to 600, 800 or 1000 nm using a plate reader (TECAN microplate reader SPARK).

The log *K*_*OW*_ was computed as the logarithm of the ratio between the concentration of photoresist molecules in the water phase after adding octanol ([resist]_*w*_) and in the octanol phase ([resist]_*o*_) before addition of octanol and afterwards. [resist]_*o*_ was computed as the difference of the concentration of photoresist molecules in the water phase before adding octanol ([resist]) and afterwards:

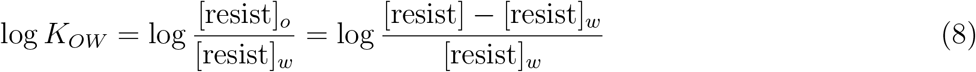

For each compound, three measurements were performed and their partition coefficients *K*_*OW*_ were calculated. Afterwards, the arithmetic mean was calculated to yield 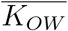 and the standard deviation as error 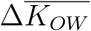. All log *K*_*OW*_ values were given as the logarithm of the averaged partition coefficient, log 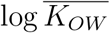, and error propagation was used to yield the error of log 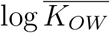 using the following formula:

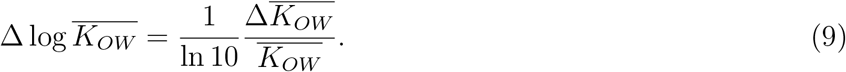

### 4.8 Electroformation of giant unilamellar lipid vesicles (GUVs)

Electroformation of GUVs was conducted in 800 mM sucrose with a a Vesicle Prep Pro device (Nanion Technologies GmbH) for all lipid mixtures. Briefly, 40 *µ*l of a 5 mM lipid mix in chloroform were spread onto the conductive side of an ITO slide. Afterwards, the slide was desiccated for 30 min. A rubber ring was placed on top of the lipids with vacuum grease and filled up with 275 *µ*l of 800 mM sucrose dissolved in water. Another ITO slide was placed on top with the conductive side facing to the bottom and the assembled chamber was connected to the electrodes of the electroformation device. The *Standard program* was chosen for generating an AC field of 3 V at 5 Hz for 2 h at 70 °C for DPPC and at 37 °C for all other lipid mixtures (DOPC, POPC, Cholesterol:DOPC in a molar ratio of 60:40, POPC:POPG in a molar ratio of 85:15). All lipid mixtures were supplemented with 0.5 mol% Atto633-DOPE for confocal imaging.

### 4.9 Fabrication of PDMS observation wells

To observe GUVs with a fluorescent microscope, PDMS based observation wells were fabricated. Briefly, PDMS elastomers and curing agent were mixed in a 10:1 weight ratio, poured into a 35 mm petri dish and degassed using a desiccator. Afterwards, the PDMS was cured in a 65 °C oven for 4 h, before cutting out 40 mm × 15 mm PDMS pieces. Several 2 mm diameter holes were then punched into the PDMS rectangles with a biopsy puncher. Afterwards, the surfaces of clean 24 mm × 60 mm glass slides and the fabricated PDMS rectangles were plasma activated for 35 s with oxygen plasma at 0.4 mbar and 200 W (PVA TePla AG), pressed together for 15 s and placed in a 65 °C oven overnight. Prior to use, the wells were coated with PVA by filling the holes with 50 g l^−1^ PVA in water solution for 5 min, before removal of the aqueous solution and blow drying with a nitrogen gun.

### 4.10 Experimental determination of permeation rates

To determine the permeation rates experimentally, GUVs in 800 mM sucrose solution were mixed in a 1:1 v/v ratio with a solution consisting of 0.1 g l^−1^ RHODB and 800 mM glucose. Right after mixing, the solution was transferred into PVA coated PDMS wells and imaged in a time series at an LSM 900 confocal laser scanning microscope (Carl Zeiss AG). For all measurements, a 20x air objective (Plan-Apochromat 20x/0.8 M27, Carl Zeiss AG) was used. Atto633-DOPE was excited with a 640 nm laser, whereas RHODB was excited with a 561 nm laser.

We performed multiple independent replica of the influx experiments (POPC N=2, POPC-POPG N=1, DOPC N=4, DOPC-Cholesterol N=2, DPPC N=2). We then ran image segmentation on the obtained confocal images with ImageJ (version 1.53c) and python 3.7 as described previously [11]. For every identified GUV *n* in the field of view at each frame of the time trace, the mean intravesicular fluorescence 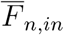, defined by the mean 8-bit grayscale pixel intensities, was divided by the mean fluorescence of the bulk medium 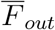:

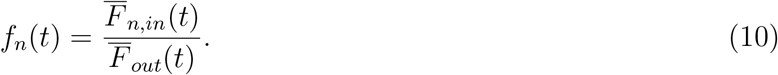

For every time point, we report the average 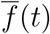 of the normalized fluorescence intensity across all GUVs in the field of view, aggregated over all time traces per condition. We report the standard deviation of the normalized fluorescence intensities as error estimate. The normalized fluorescence intensity in the GUVs follows the following relation [82]:

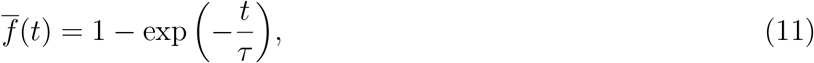

where 𝒯 represents the time point at which the mean intravesicular fluorescence reaches 0.63 times the intensity of the bulk fluorescence. We report non-linear fits of this form for the influx experiments with the square root of the estimated variance of the fitting parameter 𝒯 as error estimate. The fitting parameter 𝒯 and the radius of the GUV *R* are related to the permeation coefficient *P* [82]:

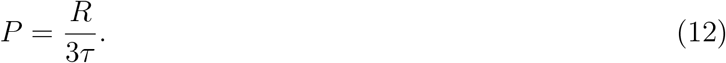

We did not derive permeability estimates from our experimental data as the intrinsically fast kinetics of the experiments did not allow an accurate determination of the fit, i.e. the normalized fluorescence of the first data point was generally greater than 0.63. To be able to relate the estimates from simulations to the experiments, we conservatively estimated the lower boundary for the permeability of Rhodamine B for the POPC GUVs. To do so, we used the time point of the first measurement 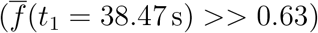 as *τ* and assumed an average GUV radius of 12.5 *µ*m [11]. Using eq. 12, we report a lower boundary of the permeation coefficient of Rhodamine B Acrylate for POPC GUVs.

## Supporting Information

Supporting Information is available from the Wiley Online Library or from the author.

## Acknowledgements

L.D., M.B., T.A., F.G., K.G., and C.A.-S. acknowledge funding from the Deutsche Forschungsgemeinschaft (DFG, German Research Foundation) under Germany’s Excellence Strategy via the Excellence Cluster 3D Matter Made to Order (EXC-2082/1 – 390761711). K.G. thanks the Hector Fellow Academy. T.A. thanks the Carl Zeiss Foundation for financial support. L.D., C.A.-S. and F.G. appreciate the financial support by the Klaus Tschira Foundation. We thank the state of Baden-Württemberg through bwHPC and the DFG through grant INST 35/1134-1 FUGG.

## Table of Contents

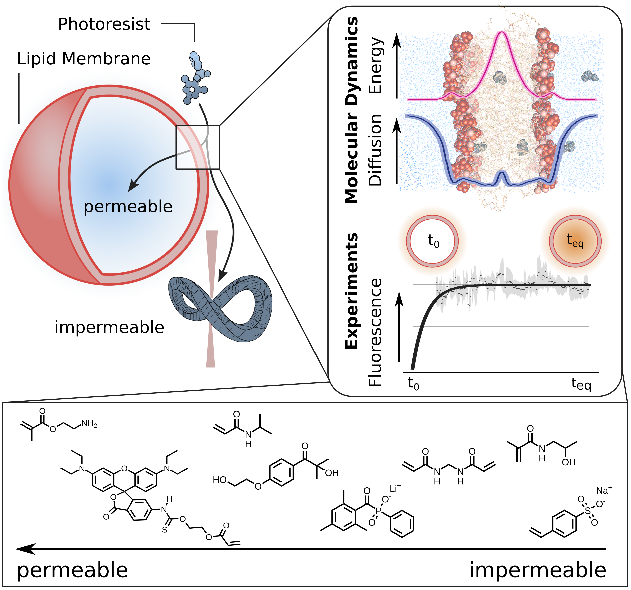

Molecular dynamics simulations and experiments are employed to elucidate the energetics and kinetics of membrane permeation of photoresist molecules that are commonly used in 3D bioprinting. This provides molecular insights into the cytocompatibility of these molecules. In the future, this will help in the rational selection of photoresists for bioprinting applications.

## Notes

### Competing Interest Statement

The authors have declared no competing interest.

