## Supporting Information for "Energetics and kinetics of membrane permeation of photoresists for bioprinting"

### Energetics of permeation of photoresists across biological lipid bilayers - Supplementary Information

#### Contents

**S1 Supplementary figures** **2**

**S2 Supplementary tables** **10**

**S3 Error estimation for permeation rates** **11**

### S1 Supplementary figures

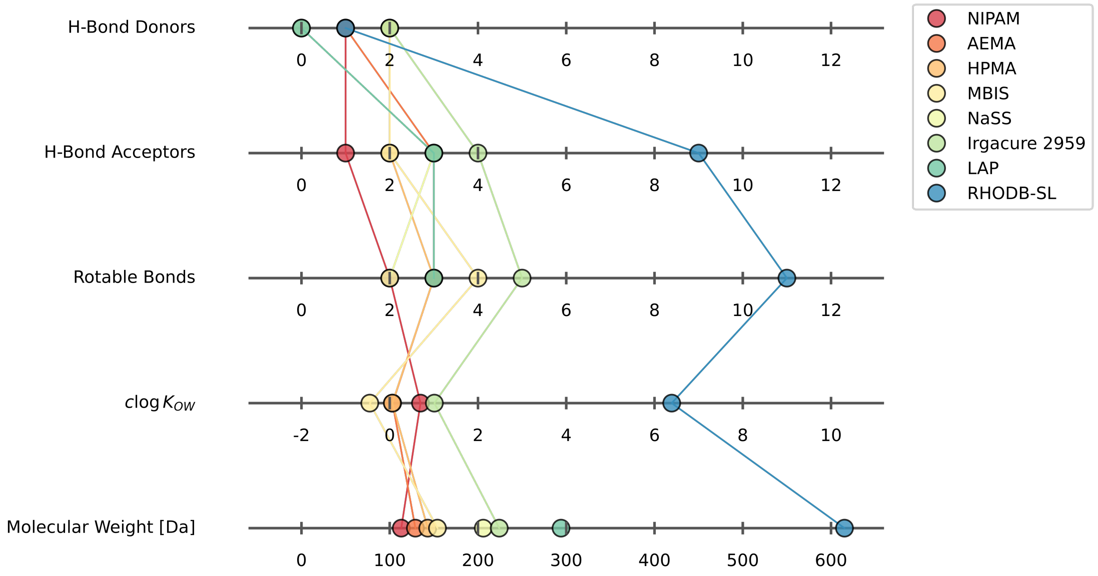


Figure S1: **Selected physico-chemical properties of the photoresists used in this study.** Depicted are five physico-chemical properties that are commonly used in pharmaceutical research to estimate the membrane permeability (respectively oral availability) of drugs [1, 2, 3]. Hence, the examined photoresist molecules span a wide range of physicochemical properties. $c \log K_{OW}$: Octanol-water partition coefficient for non-ionic molecules as predicted by RDKit [4, 5].

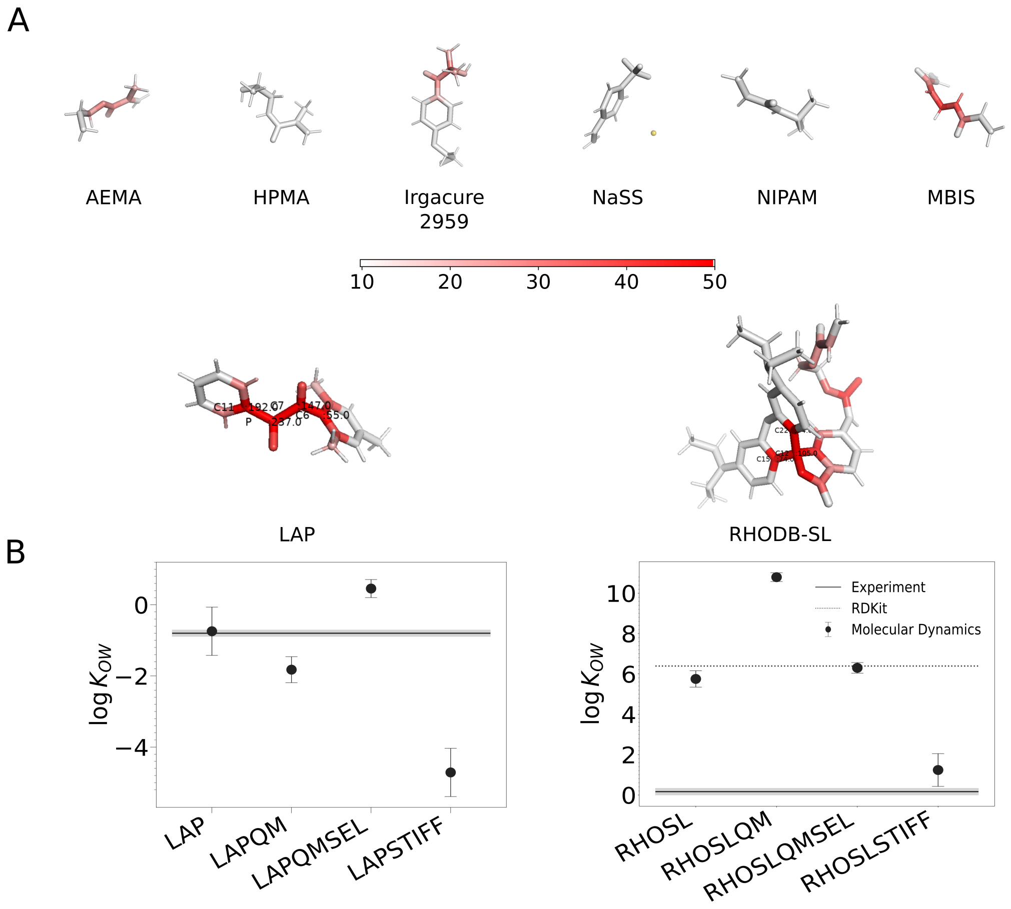


Figure S2: **Assessment of the quality of parametrization of the photoresists used in this study.** A) Heuristic error estimates of CHARMM-GUI parameterization mapped on to three dimensional structure of photoresists (Graphics: PyMOL [6]). Darker reds indicate higher errors (refer to scale bar). Generally, values below 10 correspond to confident assignment of parameters to the respective atom, while for error estimates above 50 an extra validation of the parameters is recommended. We observed errors below 40 (MBIS C4: 40.070) for all resists except for LAP and RHODB-SL where the phosphinate group (LAP) respectively spiro-moiety (RHODB-SL) were associated with a high error estimate. We tested different modifications of the bonded parameters and charges. *Molecule-QM*: Assignment of charges from DFT calculations. *Molecule-QMSEL*: Manually curated assignment of charges based on QM calculations. *Molecule-STIFF*: Modification of important dihedrals (see methods). We then compared the computed $\log K_{OW}$ estimates for these molecules with experiments and RDKit-predictions. B) Comparison of MD-derived octanol-water partition coefficients with references from experiments or machine learning predictions. For LAP, we use the experimental estimate as reference while for RHOSL, we use the prediction from RDKit [4, 5] as reference (see Results/Discussion). In both cases, because the different tested modifications did not improve the predictions, we considered the original output by CHARMM-GUI for our calculations.

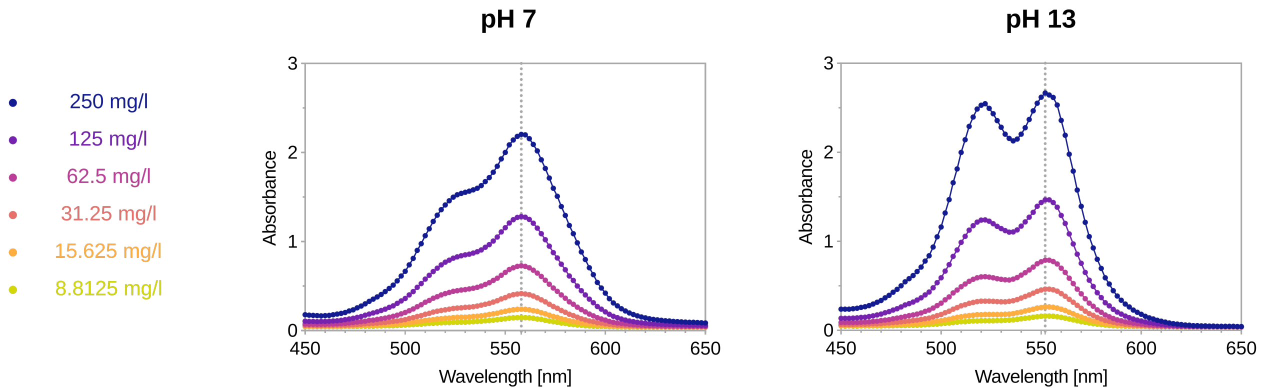


Figure S3: **Absorbance of Rhodamine B Acrylate.** Absorbance spectra of Rhodamine B Acrylate (RHODB) in water at pH=7 and pH=13 for different concentrations of the species. For determination of partition coefficients, absorbance peaks were evaluated at 558 nm and 552 nm (dotted line), respectively. In contrast to previous reports, the utilized Rhodamine dye exhibited similar spectroscopic properties at neutral and basic pH-values. This suggests that the zwitterionic form was still the dominant form in solution, since previous studies reported the spiro form of comparable Rhodamine derivatives to exhibit significantly decreased fluorescence [7, 8, 9] and absorbance [9].

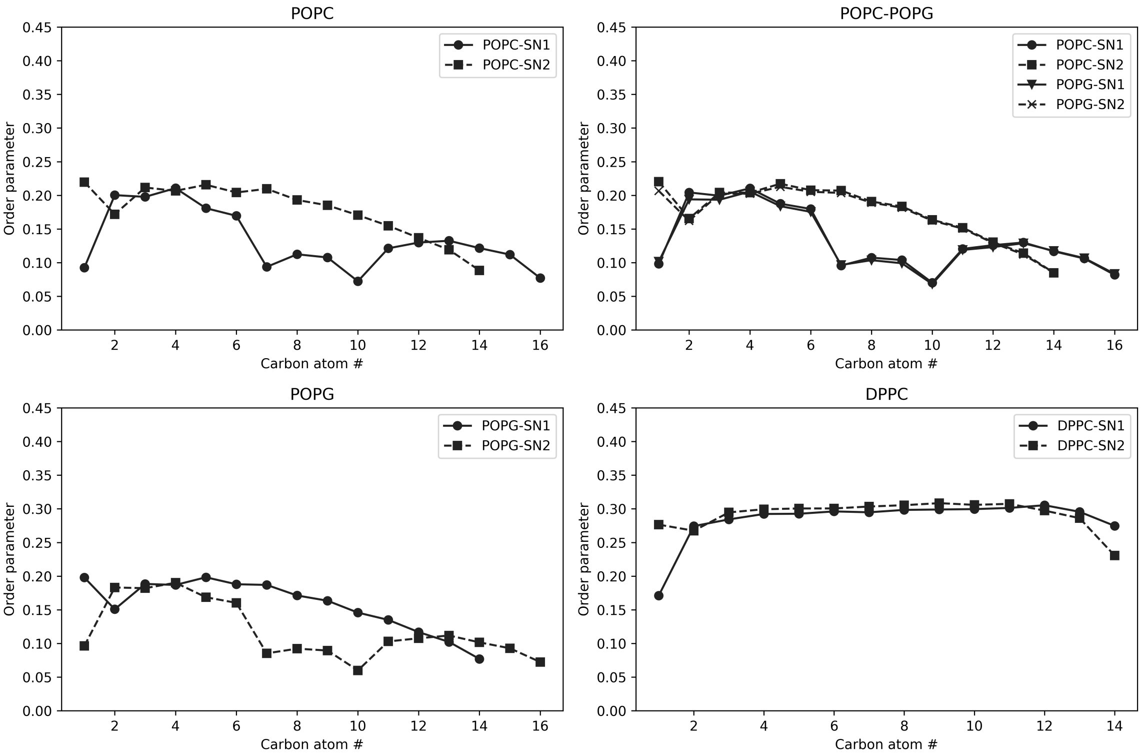


Figure S4: **Deuterium order parameter of the membranes used in this study determined with MD simulations.** Double bonds locally reduce acyl chain order, hence the drop in the local order for the SN2-lipids (monounsaturated oleoyl residues [carbons:unsaturated bonds 18:1]) for POPC, POPC-POPG and POPG membranes, compared to the SN1 lipids (saturated palmitoyl residues, [carbons:unsaturated bonds 16:0]). As expected, the different headgroups (POPC: Choline, POPG: Glycerol) do not affect the order parameters. In contrast, acyl chain order increases for highly saturated membranes like the pure DPPC membrane (here: both palmitoyl residues [carbons:unsaturated bonds 16:0]), as can be seen for the acyl chains of the DPPC membrane, simulated at 300 K. This is suggestive for the gel phase like behaviour of this membrane.

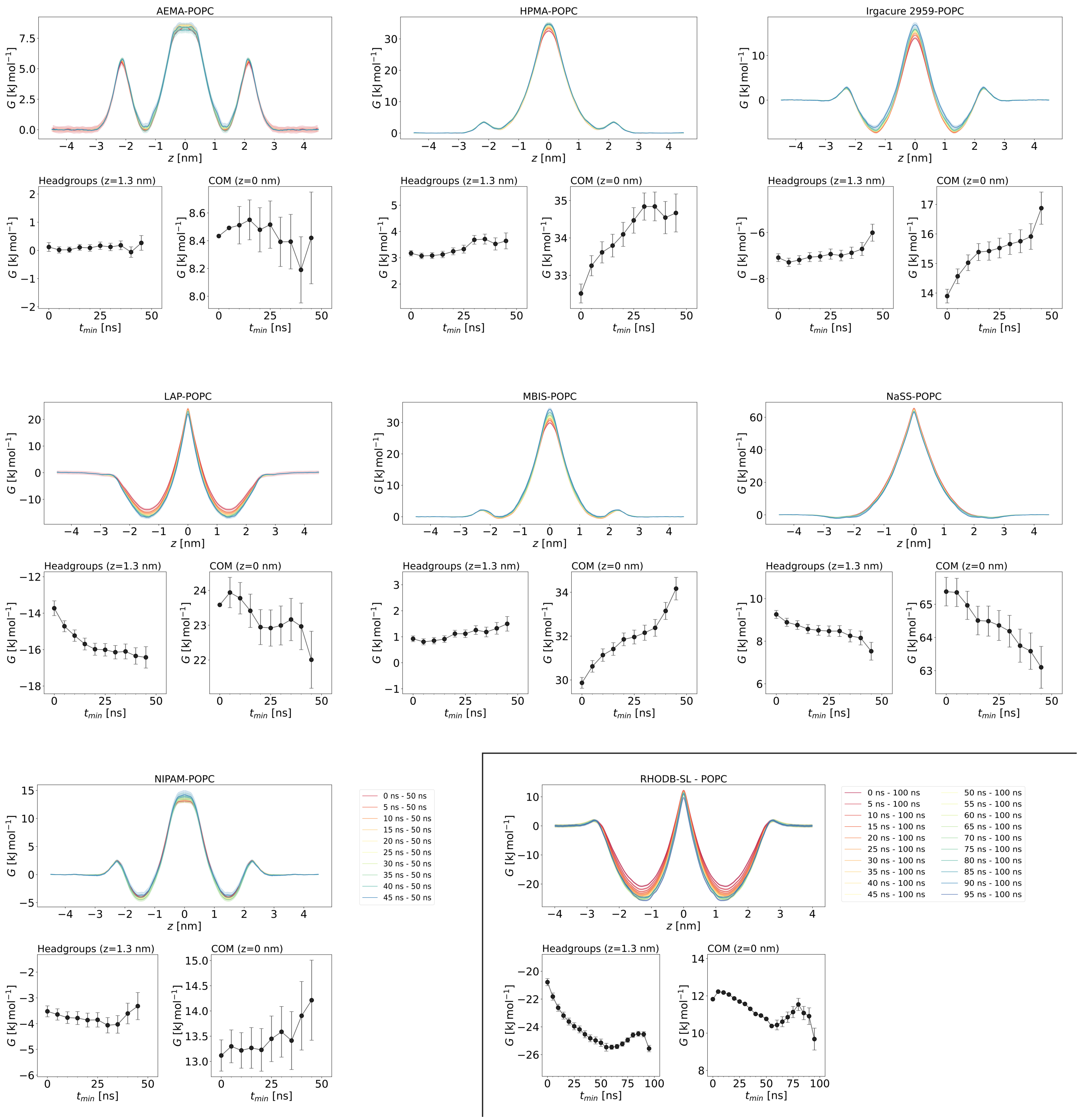


Figure S5: **Convergence of PMFs**. Blockwise backwards averaging was performed to ensure convergence of the PMFs by incrementally (in 5 ns steps) increasing the considered time for sampling with WHAM of the trajectory from 50 ns (RHODB-SL: 100 ns) backwards to $t_{min}$. Here, both free energy profiles and the exact values at the center of membrane ($z = 0$ nm) and the headgroup region ($z = 1.3$ nm) are reported. The latter indicate convergence within 50 ns for each photoresist, except RHODB-SL. For RHODB-SL, umbrella sampling simulations were extended to 100 ns to ensure convergence (*bottom, right*). The statistical uncertainty was estimated by bootstrapping and stayed below 0.5 kJ mol$^{-1}$, independent of the photoresist.

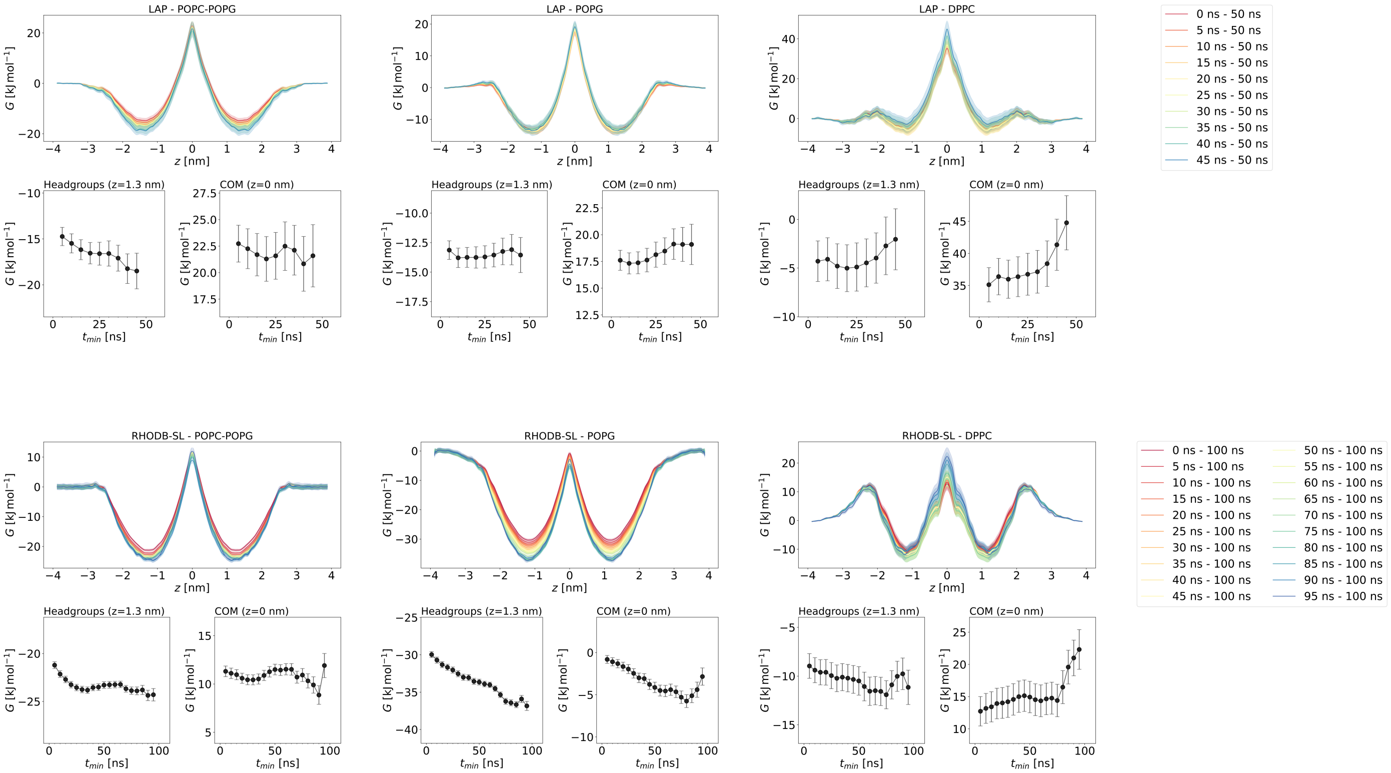


Figure S6: **Convergence of PMFs for non-POPC membranes**. Blockwise backwards averaging was performed by incrementally (in 5 ns steps) increasing the considered time for sampling with WHAM of the trajectory from 50 ns (RHODB-SL: 100 ns) backwards to $t_{min}$. Convergence of PMFs was reached within 50 ns for LAP and within 100 ns for RHODB-SL, independent of the bilayer composition.

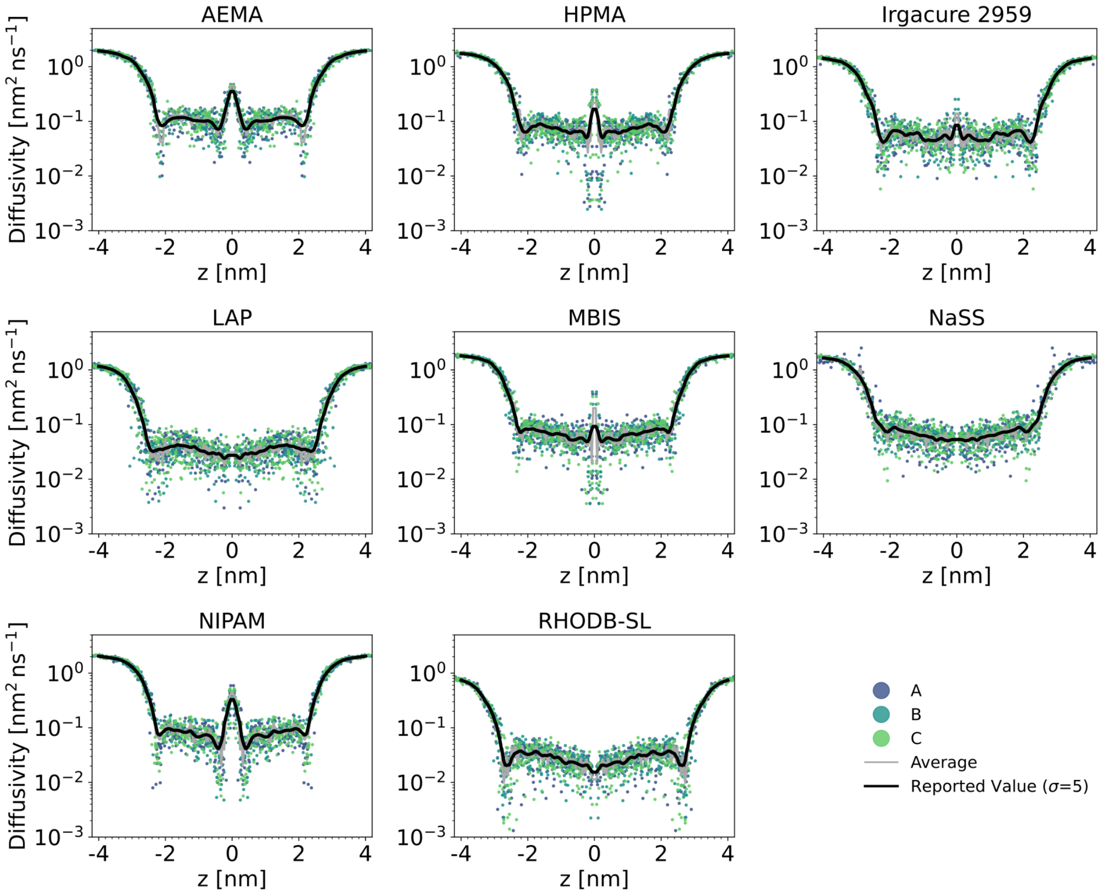


Figure S7: **Non-smoothened local diffusion coefficients for molecules of interest in POPC membrane.** Local diffusion coefficients from 3 replicates (dots), averages of the 3 replicates (gray lines) and smoothened curves (black line). For smoothing, a gaussian filter with a standard deviation of five was applied to the raw data. Smoothened curves were reported.

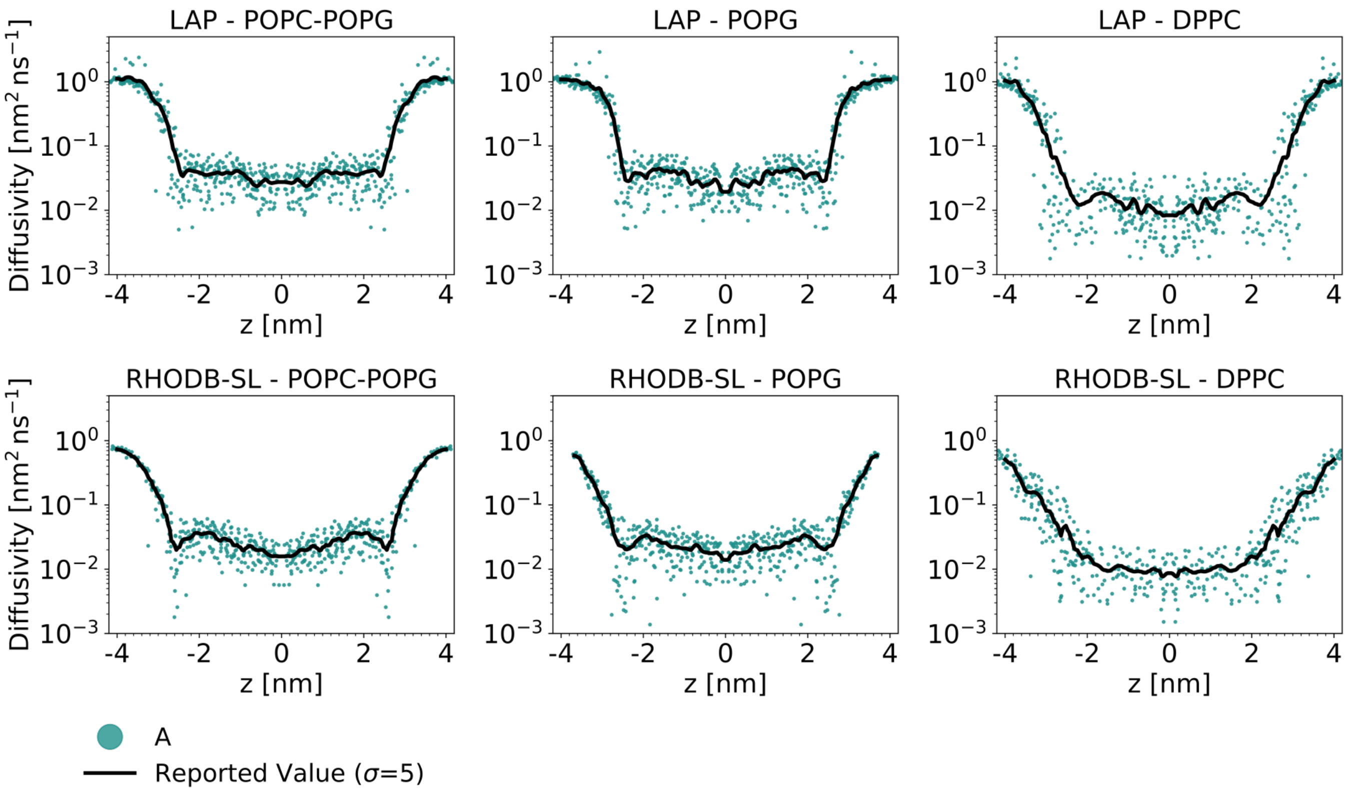


Figure S8: **Non-smoothened local diffusion coefficients for molecules of interest in POPC-POPG (85 %:15 %), POPG and DPPC membranes.** Local diffusion coefficients from one replicate (dots) and smoothened curves (black line) were used to estimate permeabilities. For smoothing, a gaussian filter with a standard deviation of five was applied to the raw data.

#### S2 Supplementary tables

Table S1: **Photoresists used in this study.** We report systematic names and aliases that were used in this study for simplicity. If applicable, we provide the PubChem ID of the compound. The provided SMILES codes were used as input on the CHARMM-GUI website [10, 11].

| Full name | Alias | PubChem ID | SMILES |
|---|---|---|---|
| N-Isopropylacrylamide | NIPAM | 16637 | CC(C)NC(=O)C=C |
| N-(2-Hydroxypropyl)methacrylamide | HPMA | 38622 | CC(CNC(=O)C(=C)C)O |
| Lithium Phenyl(2,4,6-trimethylbenzoyl)phosphinate | LAP | 68384915 | [Li+].CC1=CC(=C(C(=C1)C) C(=O)P(=O)(C2=CC=CC=C2) [O-])C |
| Sodium 4-vinylbenzenesulfonate | NaSS | 3571582 | C=CC1=CC=C(C=C1)S(=O)(=O) [O-].[Na+] |
| N-[(prop-2-enoylamino)methyl]prop-2-enamide | MBIS | 8041 | C=CC(=O)NCNC(=O)C=C |
| 2-Hydroxy-1-(4-(2-hydroxyethoxy)phenyl)-2-methylpropan-1-one | Irgacure 2959 | 86266 | CC(C)(C(=O)C1=CC=C(C=C1) OCCO)O |
| 2-aminoethyl methacrylate | AEMA | 75496 | CC(=C)C(=O)OCCN |
| Rhodamine B Acrylate (zwitterion) | RHODB-ZW | N/A | [H]N(C(=S)OCCOC(=O)C=C) C1=CC=C(C([O-])=O) C(=C1) C1=C2C=CC(C=C2OC2=C1C =CC(=C2)N(CC)CC)=[N+](CC) CC |
| Rhodamine B Acrylate (spiro-lactone) | RHODB-SL | N/A | [H]N(C(=S)OCCOC(=O)C=C) C1=CC=C2C(=O)OC3(C2=C1) C1=CC=C(C=C1OC1=C3C=CC (=C1)N(CC)CC)N(CC)CC |

Table S2: **Membranes and their composition used in this study.** Fraction as well as number of lipids per membrane in the simulations are indicated in parenthesis. * Asterisks indicate membranes that were only evaluated experimentally.

| Membrane Name | Phospholipid 1 | Phospholipid 2 | Cholesterol |
|---|---|---|---|
| POPC | POPC (100 %, 162) | - | - |
| DOPC* | DOPC (100%) | - | - |
| DPPC | DPPC (100 %, 162) | - | - |
| POPG | POPG (100 %, 162) | - | - |
| POPC-POPG | POPC (85 %, 138) | POPG (15 %, 24) | - |
| CHOL-DOPC* | DOPC (40%) | - | Cholesterol (60 %) |

Table S3: **Membrane properties determined in this study in comparison with previous studies.** All values were determined based on the last 200 ns of the production run. The area per lipid (APL) represents the average ± standard deviation of the area per lipid in this time period. Phosphate-Phosphate distances were determined by fitting two gaussians to the symmetrized and centered density profiles of the phosphoatoms in the membrane and summing up the obtained means. Temperatures between reference studies and membranes in this study may vary. Reported: Values obtained in this study. Reference: Experimental data. † Different fits to experimental data reported.

| Membrane | T [K] | APL [$nm^2$] | | | $D_{P-P}$ [nm] | | |
|---|---|---|---|---|---|---|---|
| | | Reported | Reference (T [K]) | | Reported | Reference (T [K]) | |
| POPC | 310 | $0.644 \pm 0.12$ | 0.643 | [12] (303) | 3.83 | 3.98 | [12] (303) |
| POPC 85 % - POPG 15 % | 310 | $0.646 \pm 0.012$ | - | - | 3.85 | - | - |
| POPG | 310 | $0.676 \pm 0.014$ | 0.663 | [13] (303) | 3.63 | 3.73 | [13] (303) |
| DPPC | 300 | $0.496 \pm 0.012$ | 0.472 - 0.502 † | [14] (293) | 4.40 | 4.56 - 4.84 † | [14] (293) |

#### S3 Error estimation for permeation rates

We numerically evalutated the integral in the inhomogenous solubility-diffusion (ISD) model [15] with the trapezoid rule. This can be used to derive the resistance $R$ from the previously calculated free energy and diffusion coefficient profiles, where the permeability is $P = 1/R$.

$$R = \int_{-h/2}^{h/2} \exp\left(\beta \Delta G(z)\right) \cdot D(z)^{-1} \, \mathrm{d}z, \tag{1}$$

where $h$ denotes the thickness of the membrane, $D(z)$ the diffusion coefficient, $\Delta G(z) = G(z) - G_w$ the position-dependent free energy with respect to the bulk water and the thermodynamic $\beta = 1/(k_B T)$. Numerical integration of eq. 1 was performed by discretizing the coordinate perpendicular to the membrane surface $z$ into $i = 0, ..., n$ grid points to obtain the local resistance $R(z_i)$ at each position $z_i$:

$$R(z_i) = \exp\left(\beta \Delta G(z_i)\right) \cdot D(z_i)^{-1}. \tag{2}$$

Hence, we obtained an estimate for the total resistance $R$ using the trapezoidal rule, in which the area under the curve is calculated at each position $z_i$ and then summed for all $n + 1$ grid points.

$$\int_{z_0}^{z_n} R(z) \, \mathrm{d}z \approx \sum_{i=1}^{n} \frac{R(z_{i-1}) + R(z_i)}{2} \cdot \Delta z_i, \tag{3}$$

with $\Delta z_i = z_i - z_{i-1}$, $z_0 = -h/2$, $z_n = h/2$ and $z_{i-1} < z_i$. This means that every value except the first and last will occur twice in the sum, weighted by the respective $\Delta z_i$ widths. At each grid point $z_i$, the propagated uncertainty of the local resistance $e(R(z_i))$ is composed of the error of the free energy estimate $e(G(z_i)$ and the error of the diffusivity estimate $e(D(z_i))$ according to:

$$e(R(z_i))^2 = \left( \frac{\beta \cdot \exp\left(\beta \Delta G(z_i)\right)}{D(z_i)} \cdot e(G(z_i)) \right)^2 + \left( \frac{\exp\left(\beta \Delta G(z_i)\right))}{-D(z_i)^2} \cdot e(D(z_i)) \right)^2. \tag{4}$$

Propagation of uncertainties in the sum (eq. 3) finally results in an error estimate for the resistance:

$$e(R)^2 = \left( \frac{e(R_0) \cdot \Delta z_1}{2} \right)^2 + \left( \frac{e(R_n) \cdot \Delta z_n}{2} \right)^2 + \sum_{i=1}^{n-1} \left( \frac{e(R_i) \cdot (\Delta z_i + \Delta z_{i+1})}{2} \right)^2, \tag{5}$$

with the relation $P = 1/R$, this yields the uncertainty of the permeability:

$$e(P) = e(R) \cdot R^{-2}. \tag{6}$$

Note that this represents the uncertainty of the measurement but does not account for the error introduced by the numerical evaluation of the integral.
